# NanoAmpli-Seq: A workflow for amplicon sequencing for mixed microbial communities on the nanopore sequencing platform

**DOI:** 10.1101/244517

**Authors:** Szymon T Calus, Umer Z Ijaz, Ameet J Pinto

## Abstract

**Background:** Amplicon sequencing on Illumina sequencing platforms leverages their deep sequencing and multiplexing capacity, but is limited in genetic resolution due to short read lengths. While Oxford Nanopore or Pacific Biosciences platforms overcome this limitation, their application has been limited due to higher error rates or smaller data output.

**Results:** In this study, we introduce an amplicon sequencing workflow, i.e., NanoAmpli-Seq, that builds on Intramolecular-ligated Nanopore Consensus Sequencing (INC-Seq) approach and demonstrate its application for full-length 16S rRNA gene sequencing. NanoAmpli-Seq includes vital improvements to the aforementioned protocol that reduces sample-processing time while significantly improving sequence accuracy. The developed protocol includes chopSeq software for fragmentation and read orientation correction of INC-Seq consensus reads while nanoClust algorithm was designed for read partitioning-based *de novo* clustering and within cluster consensus calling to obtain full-length 16S rRNA gene sequences.

**Conclusions:** NanoAmpli-Seq accurately estimates the diversity of tested mock communities with average sequence accuracy of 99.5% for 2D and 1D2 sequencing on the nanopore sequencing platform. Nearly all residual errors in NanoAmpli-Seq sequences originate from deletions in homopolymer regions, indicating that homopolymer aware basecalling or error correction may allow for sequencing accuracy comparable to short-read sequencing platforms.

## Background

Amplicon sequencing, particularly sequencing of the small subunit rRNA (SSU rRNA) gene and internal transcribed spacer (ITS) regions, is widely used for profiling of microbial community structure and membership [1–4]. The wide-scale application of amplicon sequencing has been driven mainly by the ability to multiplex 100’s of samples on a single sequencing run and obtain millions of sequences of target communities on high-throughput sequencing platforms [1, 4]. The primary limitation of these commonly used technologies (i.e., Illumina’s MiSeq, Ion Torrent PGM, etc.) is that their read lengths are short, ranging from 150–400 bp [5]. While excellent at bulk profiling of microbial communities through multiplexed deep sequencing, short read lengths are limited in the taxonomic resolution of sequenced reads and more so, are not amenable to robust phylogenetic analyses to assess the relationship between sequences originating from unknown microbes with those in publicly available databases. An important effect of the proliferation in short read sequencing applications has been a decrease in the rate at which long higher quality sequences, particularly of SSU rRNA genes, are being deposited in public databases. This effect is to some extent being mitigated through assembly and curation of near full-length SSU rRNA genes from metagenomic datasets [6–9] and will continue to be mitigated with novel approaches for SSU rRNA sequencing using synthetic long read approaches [10].

The introduction of long-read single molecule sequencing platforms, such as PacBio’s single-molecule real-time sequencing (SMRT) and single molecule sensing technologies on the Oxford Nanopore Technologies (ONT) MinION^TM^ platform, has opened the possibility of obtaining ultra-long reads [5, 11]. While sequencing throughput and raw data quality of long read single molecule sequencing approaches are yet to rival that of short read platforms, the ability to obtain ultra-long reads can overcome several limitations of the latter [12]. For instance, long-read sequencing combined with various error correction approaches [13, 14] has been used to obtain high-quality single contig microbial genomes [14] or increase assembly quality of previously sequenced but fragmented eukaryotic genomes assemblies [15, 16], which was not feasible using short-read approaches. Long-read sequencing capabilities have also been recently leveraged to sequence near full-length SSU rRNA genes (e.g., 16S rRNA) [17–21] or even the entire *rrn* operon [20, 22].

A majority of the studies utilizing either the SMRT or nanopore sequencing platforms have limited data their analyses efforts to sequence classification, due to the fact that widely used sequence classifiers are tolerant of high sequencing error rates [23, 24]. However, these classification-only approaches are limited in their ability to differentiate between closely related sequences, risks false detections (i.e., read incorrectly classified at the family or genus levels due to high error rates), and are unable to identify organisms that are not represented in the reference databases. In contrast, some studies have gone beyond sequence classification by using consensus sequence construction to improve overall sequence accuracy. The consensus sequence creation efforts thus far can be categorized into two approaches. The first approach involves mapping raw, noisy reads to custom or publicly available reference databases (i.e., SILVA) [25]. Subsequently, reads mapping to the same reference sequence are then used for the semi-automated or manual construction of a consensus sequence using overlapping alignments [20, 22]. While this approach does result in improved accuracy of the consensus sequence, clustering of reads based on mapping of noisy reads to reference databases has significant limitations. First, incorrect read mapping to a reference is prevalent due to high error rates of raw reads. Second, the reliance on a reference database ensures that reads originating from organisms not represented in the reference database are ignored. The more robust alternative towards high accuracy consensus sequence generation would be a completely *de novo* approach, i.e., generation of a consensus sequence without the use of any reference database.

To our knowledge, there are three reports of *de novo* data processing to reduce error rate from long read sequencing of amplicons from mixed microbial communities [17, 21, 26]. Both Singer et al and Schloss et al [17] utilized the circular consensus sequencing approach of SMRT sequencing coupled with a range of quality filtering (i.e., mismatches to primer, quality scores) and sequence clustering (i.e., pre-cluster) to generate consensus sequences from reads clustered into operational taxonomic units (OTUs) and achieved error rates of 0.5% [26] and 0.027% [17] for full-length 16S rRNA gene sequencing libraries. For the later effort [17], the number of operational taxonomic units (OTUs) of the processed data were also highly similar to the theoretical number of OTUs in tested mock communities, indicating that the application of this protocol for naturally derived mixed microbial communities is likely to result in robust diversity estimates. Li et al. [21] developed Intramolecular-ligated Nanopore Consensus Sequencing (INC-Seq) protocol for consensus-based error correction of nanopore sequencing reads with a median accuracy of 97–98%. The INC-Seq workflow involves amplicon concatermization to link multiple identical copies of the same amplicon on a single DNA molecule, sequencing of the concatamerized molecules using 2D sequencing chemistry on the nanopore sequencing platform, followed by consensus-based error correction after aligning the physically linked concatemers on each sequenced DNA strand. By using this approach, Li et al [21] were able to increase the median sequence accuracy of processed reads to 97–98%. While this significant improvement allowed for taxonomic classification of sequences to the species level, it did not allow for sequence clustering for diversity estimation due to residual median error rates of approximately 2–3%.

In this study, we leverage and expand on the INC-Seq protocol developed by Li et al [21] to provide a complete workflow for amplicon sequencing and *de novo* data processing, NanoAmpli-Seq, applied to near full-length 16S rRNA gene of mock communities that results in high-quality sequences with a mean sequence accuracy of 99.5±0.08%. The current version of NanoAmpli-Seq includes modifications to the library preparation protocol for INC-Seq and fixes a key issue with INC-Seq consensus sequences while adding a novel read partitioning-based sequence clustering approach which results in an accurate estimation of diversity of mixed microbial communities and results in higher sequence accuracy by allowing within OTU sequence alignment and consensus calling. Further, we demonstrate that NanoAmpli-Seq works equally well on the (now obsolete) 2D sequencing chemistry and the recently released 1D2 sequencing chemistry on the MinION^TM^ device. While important limitations such as suboptimal re-construction of community structure and error rate of ~0.5% remain, the proposed approach may be used for sequencing of long amplicons from complex microbial communities to assess community membership with cautious utilization of sequences from low abundance OTUs due to likely lower sequence accuracy ranging from 99–99.5% accuracy.

## Results

### Experimental design and workflow

The NanoAmpli-Seq protocol was developed and validated using amplicon pools consisting of near full-length 16S rRNA gene of a single organism (*Listeria monocytogens*) or an equimolar amplicon pool of near full-length 16S rRNA genes from 10 organisms (Supplementary Table S1). The amplicon pools were generated by PCR amplifying near full-length 16S rRNA genes from genomic DNA of the target organism(s) using primers and PCR reaction conditions as described in the materials and methods section. The respective amplicon pools were subsequently prepared for sequencing using the INC-Seq workflow as outlined in Figure 1, with a few significant modifications. Briefly, the amplicon pools were self-ligated to form plasmid-like structures which was followed by digestion with plasmid-safe DNAse to remove the remaining non-ligated linear amplicons. The DNA pool consisting of plasmid-like structures was subject to rolling circle amplification (RCA) using random hexamer-free protocol using a combination of primase/polymerase (PrimPol) and hi-fidelity Phi29 DNA polymerase [27]. The RCA product was subject to two rounds of T7 endonuclease I debranching and g-TUBE fragmentation followed by gap filling and DNA damage repair. Description of the protocol including reagent volumes and incubation conditions is provided in the materials and methods and a step-by-step protocol is provided in the supplementary text. The prepared amplicon pools for both single organism and 10 organism mock community samples were then subject to library preparation using the standard 2D (SQK-LSK208) (Runs 1 and 2) and 1D2 (SQK-LSK308) (Runs 3 and 4) kits using ONT specifications and sequenced on the MinION^TM^ MK1b device followed by basecalling using Albacore 1.2.4.

**Figure 1:**
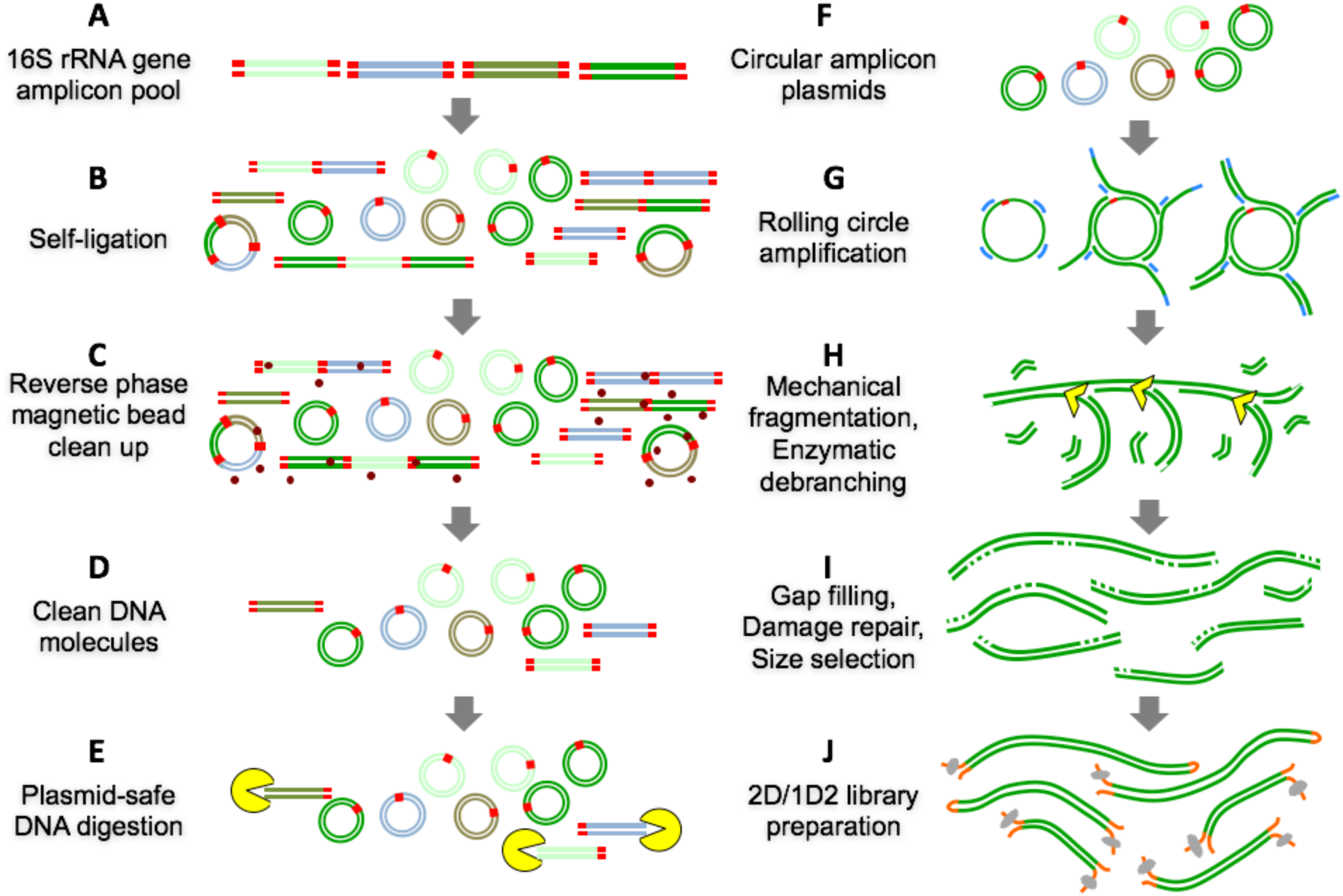
Overview of the sample preparation protocol for 16S rRNA gene pool preparation, plasmid-like structure preparation, enzymatic debranching and mechanical fragmentation, and 2D and 1D2 library preparation including intermediate clean-up steps.

Each resulting read consisted of multiple concatamerized physically linked amplicons from the one original 16S rRNA gene amplicon. The long concamtermized amplicon reads were subject to INC-Seq’s anchor based alignment and consensus error correction using three different alignment options (i.e., blastn, Graphmap, and partial order alignment (POA)) and followed by iteratively running PBDAGCON on the consensus for error correction (INC-Seq flag “iterative”). Reads with irregular segment length, unmappable anchors, and potentially chimeric molecules (i.e., concatemers from more than one original 16S rRNA gene amplicon) were removed during the generation of the INC-Seq consensus read. Manual inspection of INC-Seq consensus reads revealed that a vast majority had an incorrect orientation of primers (Figure 2A). Specifically, the forward and reverse primers did not occur at the ends of the INC-Seq consensus reads but rather were co-located at varying positions along the length of each read. Efforts to manually split INC-Seq reads and re-orient the forward and reverse splits based on primer orientation revealed the presence of tandem repeats of nearly identical sequences, which affected efforts to merge the forward and reverse read splits (Figure 2B, 2C).

**Figure 2: (A).**
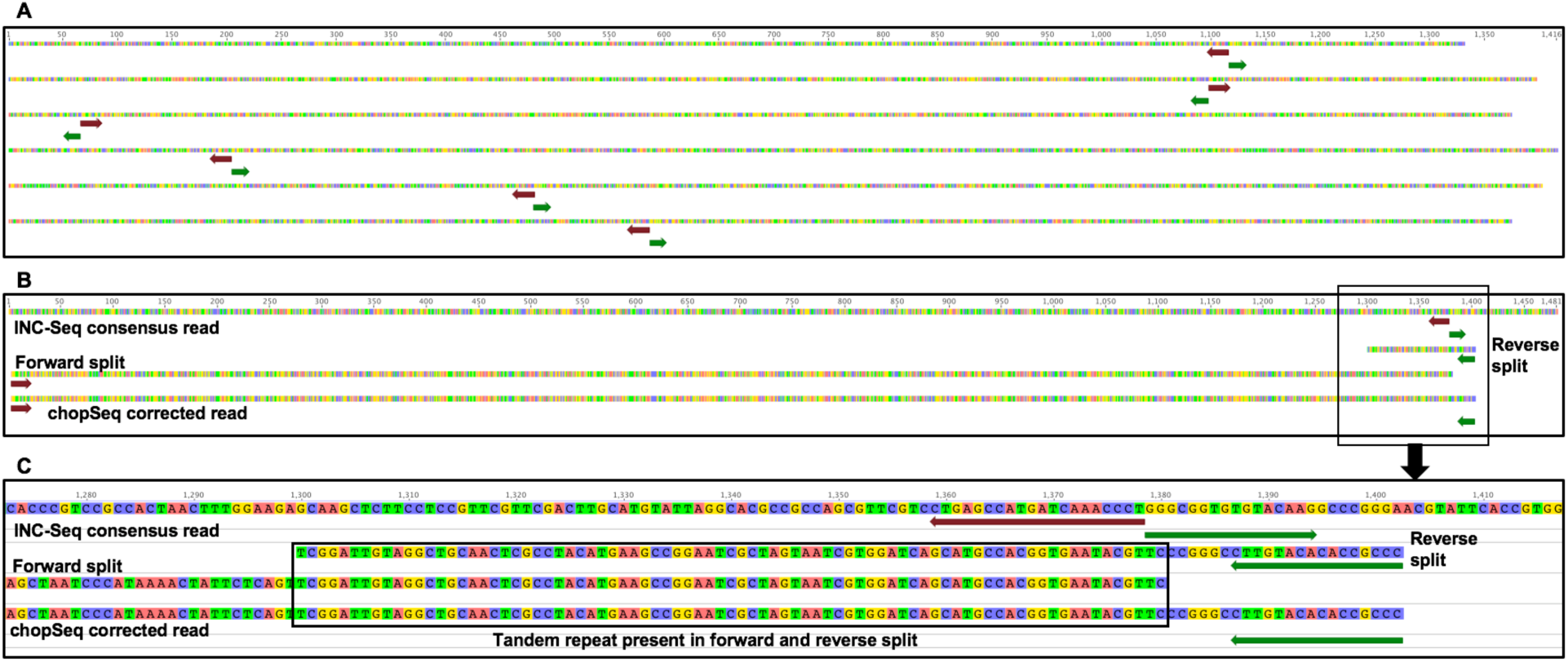
Example of INC-Seq consensus reads showing the improper orientation with forward (maroon) and reverse (green) primers co-located and incorrectly oriented. **(B)** Manual splitting and re-orientation of the reads revealed the presence of tandem repeats in the forward and reverse splits which were identified and removed using chopSeq. **(C)** An expanded view of tandem repeat region in Figure 2B.

To this end, we developed the chopSeq algorithm as part of the NanoAmpli-Seq workflow. The chopSeq algorithm uses pairwise2 open source library from Biopython package to identify user provided primers (forward “–f”, reverse “–r”) sequences including degenerate bases in the INC-Seq consensus reads. Primer detection is carried out in different orientations and primer match scores for each orientation are generated. Subsequently, primer sequences in the INC-Seq consensus read with the highest mean score are re-oriented and any overhang is removed. Re- orienting reads using primer orientation resulted in the identification of insertions consisting of repeated sequence patterns, i.e., tandem repeats. These tandem repeats were identified using etandem algorithm from EMBOS open source software package (http://emboss.bioinformatics.nl/cgi-bin/emboss/etandem) and various features of these repeats were delineated, i.e. tandem minimum repeat, tandem maximum repeat, and mismatch rate. The percent identify between tandem repeat is estimated iteratively measuring the sequence similarity between co-occurring segments using window size ranging from 10 bp to 350 bp with diminishing sequence similarity threshold with increasing window size. The sequence similarity threshold with increasing window size was applied as longer tandem repeats tend to have lower similarity to each other compared to shorter. After completing re-orientation of reads in the removal of tandem repeats, the forward and reverse splits are merged into a single read, and any read that does not match prescribed length threshold (i.e., 1300–1450 bp) is discarded. This process of primer identification and tandem repeat removal can also be visualized by turning on verbosity mode (flag = -v) and the results can be exported in fasta format.

To enable fully reference-free analyses, we developed the nanoClust algorithm which takes the fasta file of chopSeq corrected reads as input and then performs read-partitioning based *de novo* clustering using VSEARCH [28] to delineate OTUs at a user-specified sequence similarity threshold (i.e., 97% in this study) followed by within OTU read alignment and consensus calling for each OTU. The nanoClust algorithm is written in python and requires Biopython packages such as Seq, SeqIO, AlignIO, aligninfo and pairwise2. This algorithm was explicitly designed for *de novo* clustering because standard *de novo* clustering approaches such as VSEARCH [28] and the clustering approaches available in mothur [29, 30] vastly overestimated the richness of the mock community when using chopSeq corrected reads (see details below). The nanoClust algorithm takes chopSeq corrected reads in fasta format, splits the reads into partitions based on user-defined partition size, implements VSEARCH [28] for dereplication, chimera detection and removal in each partition, and clustering for each partition to identify the partition category with optimal (i.e., maximum) number of OTUs (not counting singleton OTUs) and (3) discards singletons. Following this, nanoClust extracts read IDs for each OTU bin from the best performing partition, the extracted read IDs for each OTU bin are then used to obtain full-length chopSeq corrected reads, a subset of reads that fall within 10% of the average full-length read distribution within each OTU bin are aligned using Multiple Alignment using Fast Fourier Transform (MAFFT) [31, 32] with G-INS-i option, followed by consensus calling to obtain full-length representative sequence for each OTU. The entire data processing workflow is shown in Figure 3.

**Figure 3: (A).**
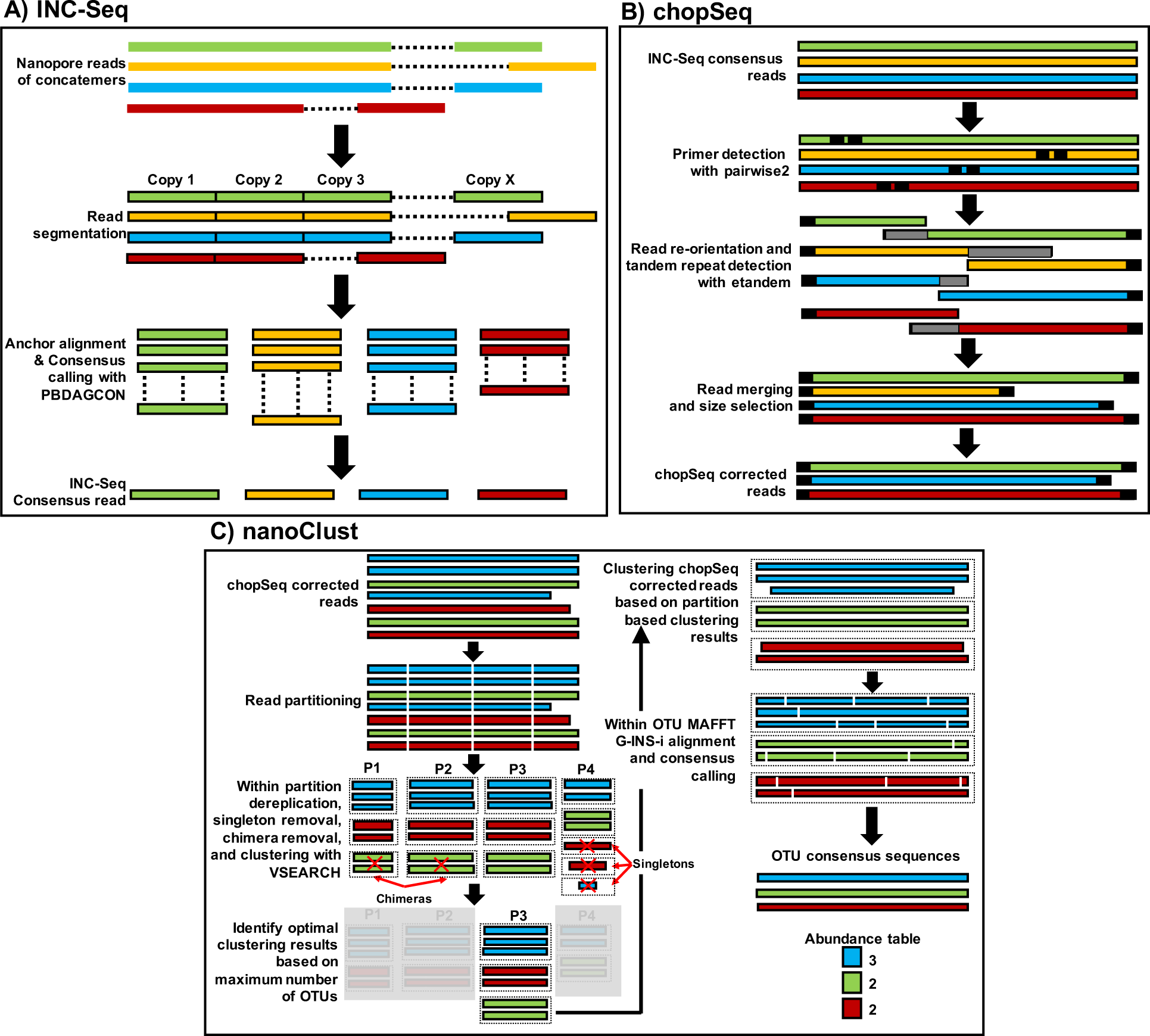
Overview of the INC-Seq based anchor alignment and iterative consensus calling using PBDAGCON. **(B)** INC-Seq consensus reads were subject to chopSeq based read reorientation followed by tandem repeat removal and size selection to retain reads between 1300-1450bp. **(C)** chopSeq corrected reads are subject to partitioning followed by VSEARCH based binning to identify optimal binning results using partition that generates maximum number of OTUs (without singletons). MAFTT-G-INS-i was then used for sequence alignment of a subset of full length reads from each OTU bin for the best performing partition and the alignment was used to create the OTU consensus read.

### Modifications to the original INC-Seq protocol significantly reduces time required for amplicon concatemer pool preparation

While the proposed DNA preparation protocol is based on the previously developed INC-Seq approach [21], it contains multiple improvements that allow for faster and more efficient library preparation. These modifications include reduced incubation times for self-ligation step and plasmid-safe DNAse digestion process. More importantly, the current protocol utilizes Tth PrimPol [27] and Phi29 DNA polymerase enzymes for RCA which minimizes the formation of unspecific products that may occur when using random hexamers. Similarly, the NanoAmpli-Seq protocol utilizes T7 endonuclease I enzyme for enzymatic debranching of RCA product combined with mechanical fragmentation step involving use of the g-TUBE. Thus, while our protocol increases the number of intermediate steps for sample preparation, by optimizing each step, it reduces the overall time required for sample DNA preparation to 6 hours (approximately 70% reduction compared to the original INC-Seq protocol). These improvements not only result in analyses of near full-length 16S rRNA gene (i.e., twice the amplicon size of the previously developed INC-Seq approach), but the combination of the improved protocol with appropriate data processing modifications resulted in significant increase in high-quality data post-processing.

### NanoAmpli-Seq data yield for 2D and 1D2 experiments

Runs 1, 2, 3, and 4 resulted in 29420, 59490, 142233, and 301432 raw records with post basecalling read lengths ranging from 5bp to 43kbp and 5bp to 234kbp for 2D and 1D2 sequencing protocols (Supplementary Figure S1). The pass reads to total raw reads ratio ranged from 28% for 2D and 7–9% for the 1D2 experiments (Table 1). It is unclear if the low yield of pass reads, particularly for the 1D2 experiments, were due to the concatemerization process, DNA damage during enzymatic debranching and mechanical fragmentation that was unrepaired in the subsequent steps, or due to basecalling issues. All pass reads were subjected to INC-Seq processing to allow for consensus-based error correction using reads with a minimum of three concatemers per read (i.e., reads with less than three concatemers were excluded for any subsequent analyses) as compared to six concatemer threshold used by Li et al [21]. The number of concatemers per read passing INC-Seq threshold ranged from 3 to 21 and 3 to 42 for 2D and 1D2 data (Supplementary Figure S1). The total number of reads passing the three concatemer threshold ranged from 36–75% of the base called reads depending on the experiment, sequencing protocol, and alignment approach during INC-Seq processing (Table 1). This was significantly higher than those reported by Li et al [21], primarily due to the use of three compared to six concatemer threshold recommended previously.

**Table 1:** Summary of the total number of reads and median read lengths at each step of the data processing workflow for all experiments.

|  | Number of Reads |  |  |  | Median Read Length |  |  |  |
| --- | --- | --- | --- | --- | --- | --- | --- | --- |
| Protocol | 2D | 1D2 | 2D | 1D2 | 2D | 1D2 | 2D | 1D2 |
| Experiment | One organism | One organism | Ten organism | Ten organism | One organism | One organism | Ten organism | Ten organism |
| Run | 1 | 4 | 2 | 3 | 1 | 4 | 2 | 3 |
| Raw records | 29420 | 301432 | 59490 | 142233 | - | - | - | - |
| Pass reads | 8108 | 20888 | 16403 | 12011 | 4868 | 7305 | 5401 | 6418 |
| INC-Seq aligner | Blastn |  |  |  |  |  |  |  |
| INC-Seq | 3911 | 7618 | 7011 | 9081 | 1394 | 1433 | 1396 | 1413 |
| chopSeq | 3911 | 7618 | 7011 | 9081 | 1383 | 1387 | 1377 | 1374 |
| chopSeq-size select | 3748 | 7288 | 5186 | 7265 | 1384 | 1388 | 1377 | 1375 |
| nanoClust | 2997 | 6153 | 3677 | 5465 | 1396 | 1398 | 1384 | 1386 |
| INC-Seq aligner | Graphmap |  |  |  |  |  |  |  |
| INC-Seq | 3902 | 7631 | 7004 | 9169 | 1400 | 1439 | 1401 | 1420 |
| chopSeq | 3902 | 7631 | 7004 | 9169 | 1384 | 1388 | 1378 | 1377 |
| chopSeq-size select | 3765 | 7399 | 5190 | 7496 | 1384 | 1389 | 1377 | 1377 |
| nanoClust | 2981 | 6179 | 4141 | 5490 | 1397 | 1396 | 1384 | 1386 |
| INC-Seq aligner | POA |  |  |  |  |  |  |  |
| INC-Seq | 3913 | 7643 | 7025 | 9191 | 1414 | 1457 | 1415 | 1443 |
| chopSeq | 3913 | 7643 | 7025 | 9191 | 1394 | 1396 | 1386 | 1387 |
| chopSeq-size select | 3779 | 7088 | 5076 | 6954 | 1394 | 1396 | 1385 | 1386 |
| nanoClust | 3184 | 5993 | 3916 | 5622 | 1398 | 1389 | 1384 | 1386 |

### INC-Seq processed reads demonstrated incorrect read orientation and presence of tandem repeats

While the median read lengths for post INC-Seq were generally in the expected range (i.e., 1350–1450 bp) (Table 1) and similar for all three alignment methods used (Supplementary Figure 2), manual inspection of the reads revealed several instances of incorrect read orientation (Figure 2). The amplicon pool preparation protocol relies on RCA of plasmid-like structures constructed through self-ligation of linear amplicons followed by a combination of enzymatic debranching and mechanical fragmentation to generate linear molecules with multiple concatemers. Considering the fragmentation and debranching steps are not driven by sequence specificity, it would be reasonable to assume that the resulting linear amplicon is unlikely to have the correct orientation, i.e. 16S rRNA gene-specific forward and reverse primers do not flank the entire amplified region. Indeed, we found a vast majority of the 2D and 1D2 INC-Seq consensus reads were incorrectly oriented for the single organism sequencing runs, with forward and reverse primers not located at the ends of the reads. As a result, the reads were chopped at the primer sites and re-oriented to allow for the forward and reverse primers to be correctly oriented. However, during the process of read re-orientation, we also discovered the presence of inserts in the form of tandem repeats. Additional inspection of these inserts revealed that they were composed of multiple repetitive sequences, with the length of these inserts ranging from 10 bp to in excess of 1500 bp (for rare cases) with median tandem repeat size ranging from 12 to 62 bp. The proportion of INC-Seq processed reads with tandem repeats varied from 60–75% but did not reveal any significant effect of type of aligner used during INC-Seq consensus calling or the sequencing chemistry itself. Interestingly, however, the length distribution for the tandem repeats was strongly associated with the sequencing chemistry (Figure 4).

**Figure 4:**
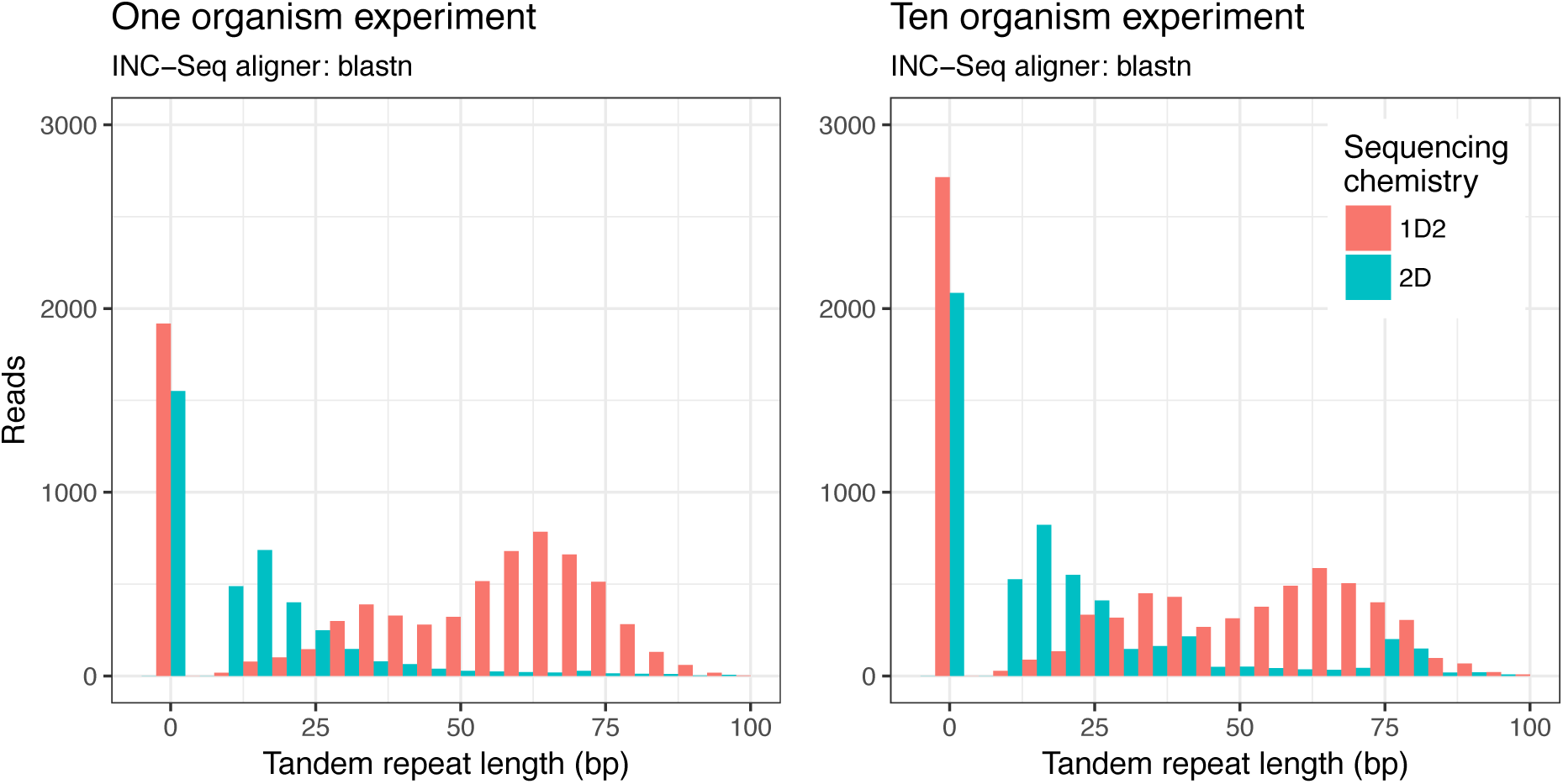
Histogram of tandem repeat length distribution of the INC-Seq processed reads did not show any effect of the aligner used during the INC-Seq process, but rather a marked effect of the sequencing chemistry. Results are only shown for INC-Seq consensus reads generated using blastn aligner.

Specifically, the 1D2 reads had longer tandem repeats as compared to the 2D reads and demonstrated a bimodal distribution of tandem repeat lengths as compared to the 2D data which showed a unimodal tandem repeat length distribution (Figure 4). While the template and complements in the 2D sequencing chemistry are physically linked by a hairpin adapter, they are not physically linked in the 1D2 sequencing chemistry; this could likely be the cause of differences in tandem repeat length distribution between 2D and 1D2 experiments.

### Read re-orientation and tandem repeat removal significantly improves sequence quality

BLASTn analyses of INC-Seq reads against reference database composed of 16S rRNA gene sequences of one (Run1 and Run4), or ten organisms (Run 2 and 3) revealed that a combination of incorrect read orientation and presence of tandem repeats significantly affected overall sequence quality. While the average sequence similarity between INC-Seq consensus reads and the reference sequence was 97±0.37%, the portion of the INC-Seq consensus read demonstrating a contiguous alignment to the reference sequence varied significantly (Figure 5); the remaining section of the read typically resulted in shorter secondary alignments with similar sequence similarity to that of the primary alignment. However, post chopSeq the average proportion of the read aligning to the reference sequences increased to 96±2.3% with additional step of discarding reads less than 1300 bp and greater than 1450 bp increasing the average proportion of the read aligned to 98.4±0.7% while the sequence similarity between chopSeq (97.5±0.42%) and chopSeq followed by size selection (chopSeq_SS) (98±0.23%) remained similar to or slightly better than INC-Seq processed read. This demonstrates that read reorientation and tandem repeat removal resulted in reconstruction of reads with a high level of similarity to the reference sequences (Figures 5).

**Figure 5:**
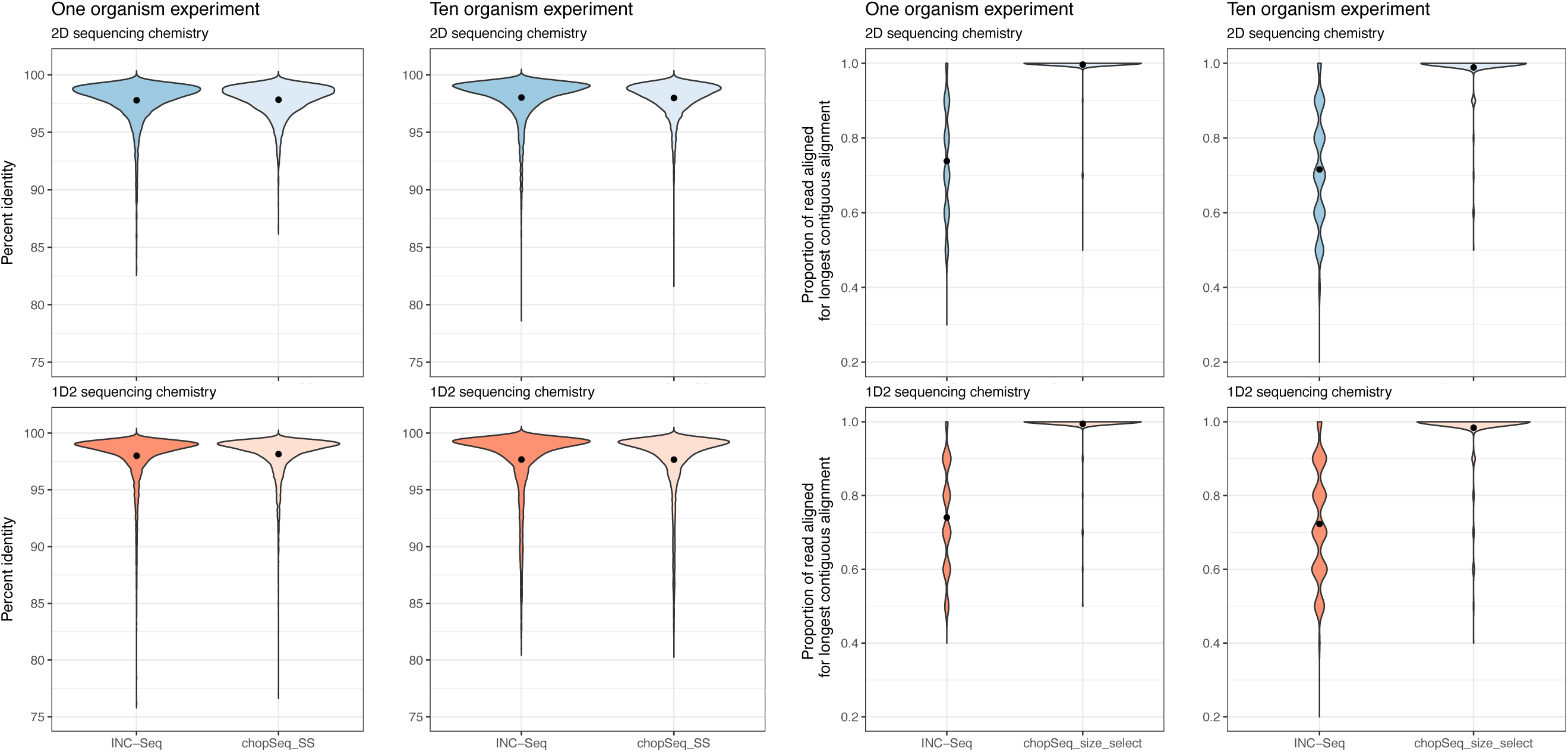
While the distribution of percent identities of INC-Seq and chopSeq processed reads (chopSeq_SS) to reference sequences was on average 97–98%, variable lengths of the INC-Seq processed reads aligned to the reference sequences. In contrast, nearly the entire length of the chopSeq processed reads aligned to the reference sequence without affecting overall sequence similarity. Results are only shown for INC-Seq reads generated using blastn aligner.

Inspections of the read to reference alignment length ratio indicated that the primary source of sequence error for both INC-Seq and chopSeq corrected reads originated from deletions; i.e., the majority of reads had a read to reference alignment ratio lower than 1. While deletions in reads were also strongly associated with sequence accuracy for post chopSeq and size selected reads, a small proportion of chopSeq corrected reads showed read to reference alignment ratios of greater than 1 (Figure 6), suggesting that insertions were less prominent than deletions.

**Figure 6:**
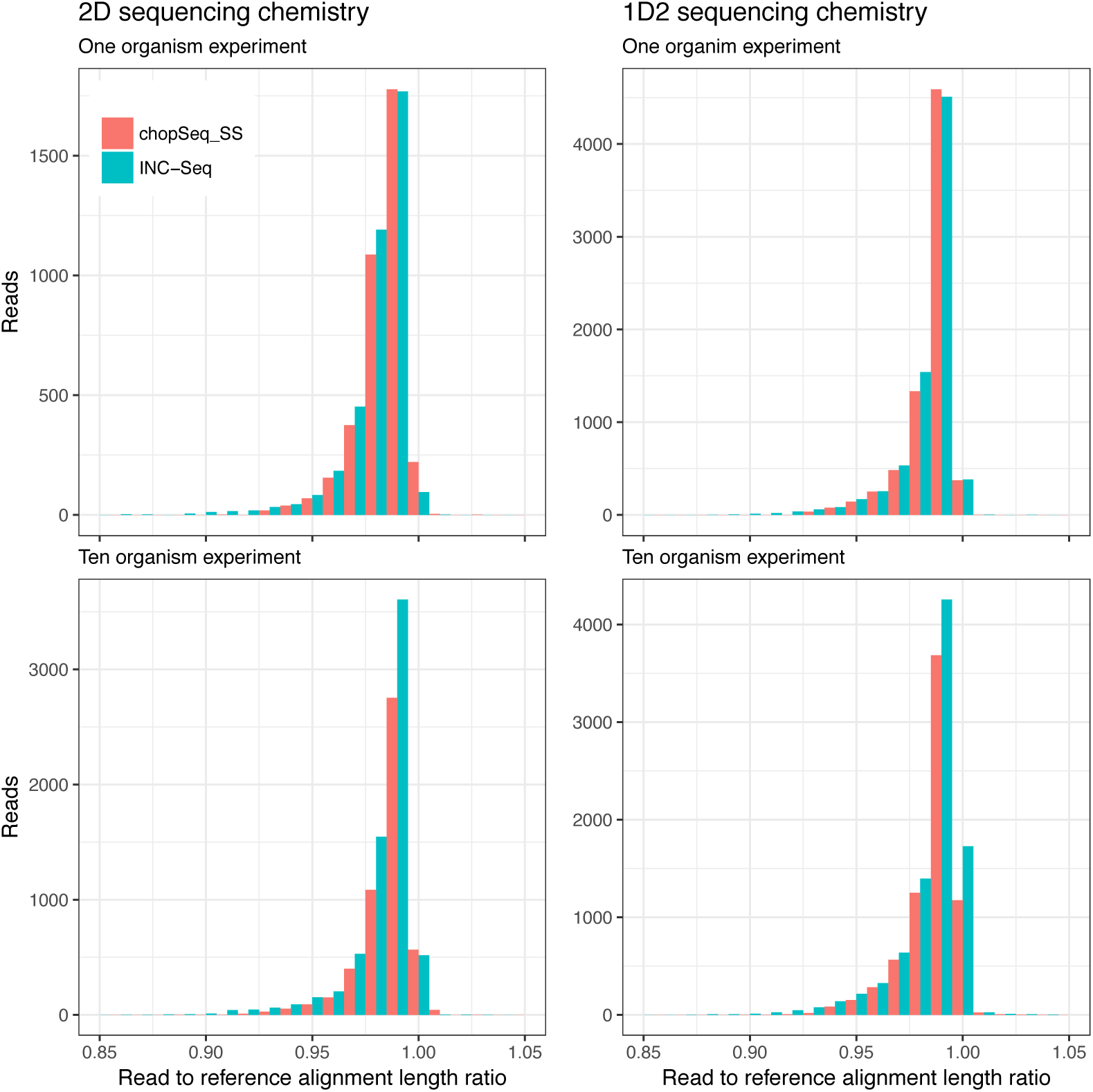
The ratio of the alignment length of INC-Seq and chopSeq corrected reads to that of the corresponding reference sequences was consistently lower than 1, suggesting that deletions in the INC-Seq and chopSeq corrected reads were the primary cause of dissimilarity with the reference sequences. Results are only shown for INC-Seq reads generated using blastn aligner.

### *De novo* clustering of chopSeq corrected sequences followed by within cluster consensus calling significantly enhances sequence accuracy

The overall sequence accuracy increased to an average of 97.9±0.23% following chopSeq read correction and size selection with 98±0.23% of the read aligning to the reference (Figure 5). However, approximately 5 and 10% of reads for the 2D and 1D2 runs, respectively, exhibited sequence accuracy of less than 95% with some sequences aligning over less than 50% of the read length even after chopSeq correction and size selection. These poor-quality reads could not be selectively filtered out based on any commonly used quality filtering criteria (e.g., maximum homopolymers length, primer mismatches, etc.) and significantly affected clustering of reads into OTUs. For instance, VSEARCH based clustering of full length post chopSeq and size selected reads (INC-Seq aligner: blastn) at a 97% sequence similarity threshold resulted in 817 (with 777 singletons) and 1301 (with 1238 singletons) for 2D and 1D2 data for single organism experiment and 2122 (with 1742 singletons) and 2725 (with 2447 singletons) for 2D and 1D2 data for 10 organism experiments. We hypothesized that accrual of residual errors over the entire read length hampered the accuracy of the OTU clustering and that accurate clustering was more likely over shorter regions of the reads due to fewer absolute errors. To this end, we developed nanoClust which utilizes partitioning of reads in user-defined lengths, followed by application of VSEARCH within each partition for dereplication, singleton removal, chimera detection and removal, clustering at user-defined sequence similarity threshold (i.e., 97% sequence similarity in this study), followed by within cluster sequence clustering and consensus calling.

We tested the effect of the choice of partition length on the estimation of the richness of the mock communities (i.e., number of observed OTUs) and overall sequence accuracy post within-OUT MAFFT-G-INS-i alignment and consensus sequence construction. To this effect, we varied the number of partitions from one (i.e., partition length of 1300 bp) to seven (i.e., partition length 180 bp). With increasing number of partitions (i.e., decreasing partition length), the number of OTUs being detected were significantly inflated above the theoretical threshold while at the same time the average sequence accuracy decreased (Figure 7).

**Figure 7:**
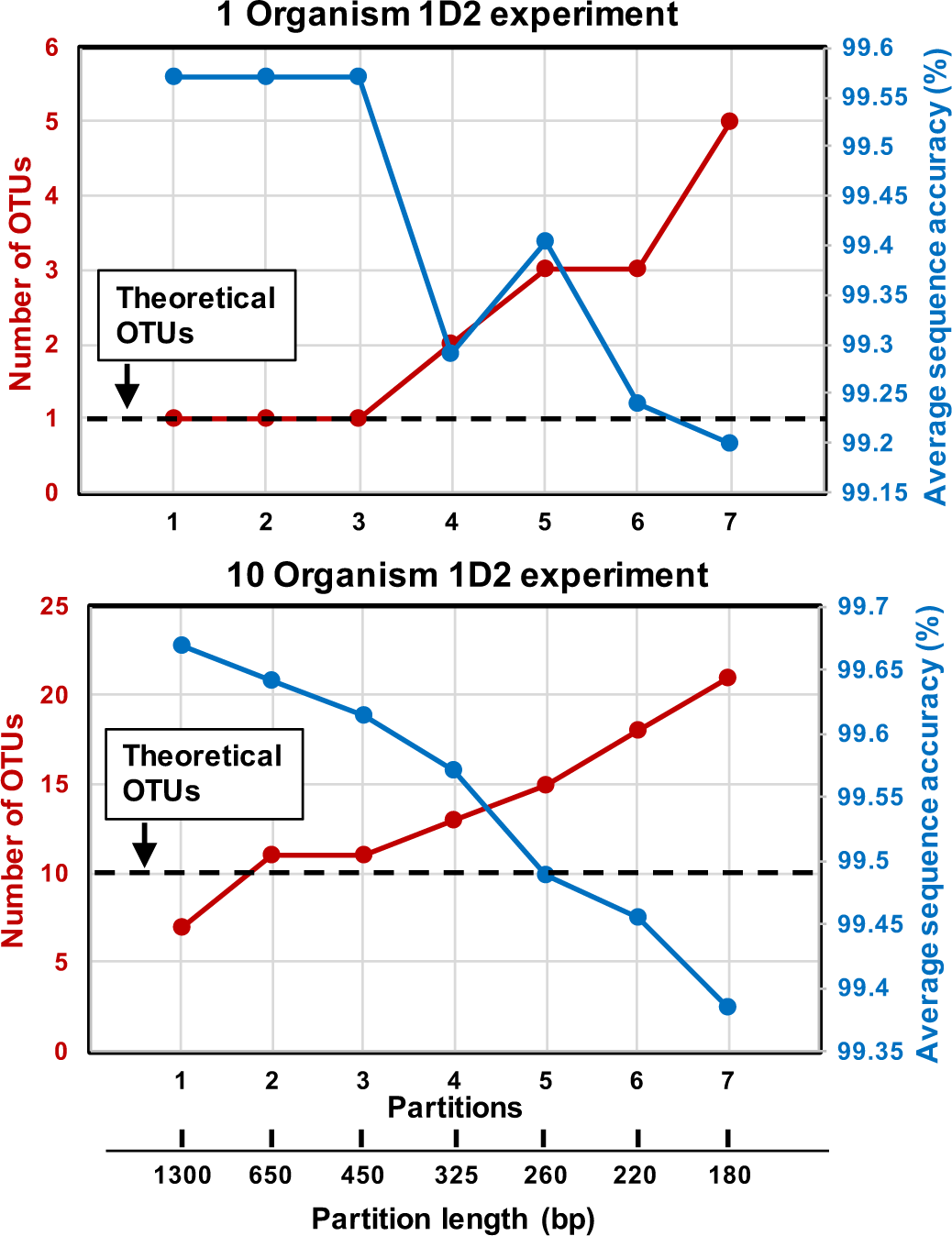
Increasing the number of partitions (and decreasing partition length) during nanoClust processing results in inflation in the number of OTUs observed and a decrease in overall sequence accuracy.

This was consistent for both the one organism and 10 organism experiments for both 1D2 and 2D experiments (Figure 7). As the partition size decreased the number of OTUs decreased and the overall sequence accuracy increased. The highest average sequence was observed with a single partition, but the number of OTUs was lower than theoretical for the 10-organism experiment. Further, using a single partition approach also resulted in discarding a significant number of sequences that were deemed singletons prior to OTU clustering. Specifically, while three less OTUs were detected in the single partition approach the total number of sequences retained post clustering were 20% lower as compared to when two or three partition approach was used. Considering the tradeoff between sequencing depth (and the resultant impact on detection of lower abundance OTUs), the extent of deviation from the theoretical number of OTUs and overall sequence accuracy, we recommend using either the two or three partition approach which result in similar outcomes on all three metrics.

**Table 2:**
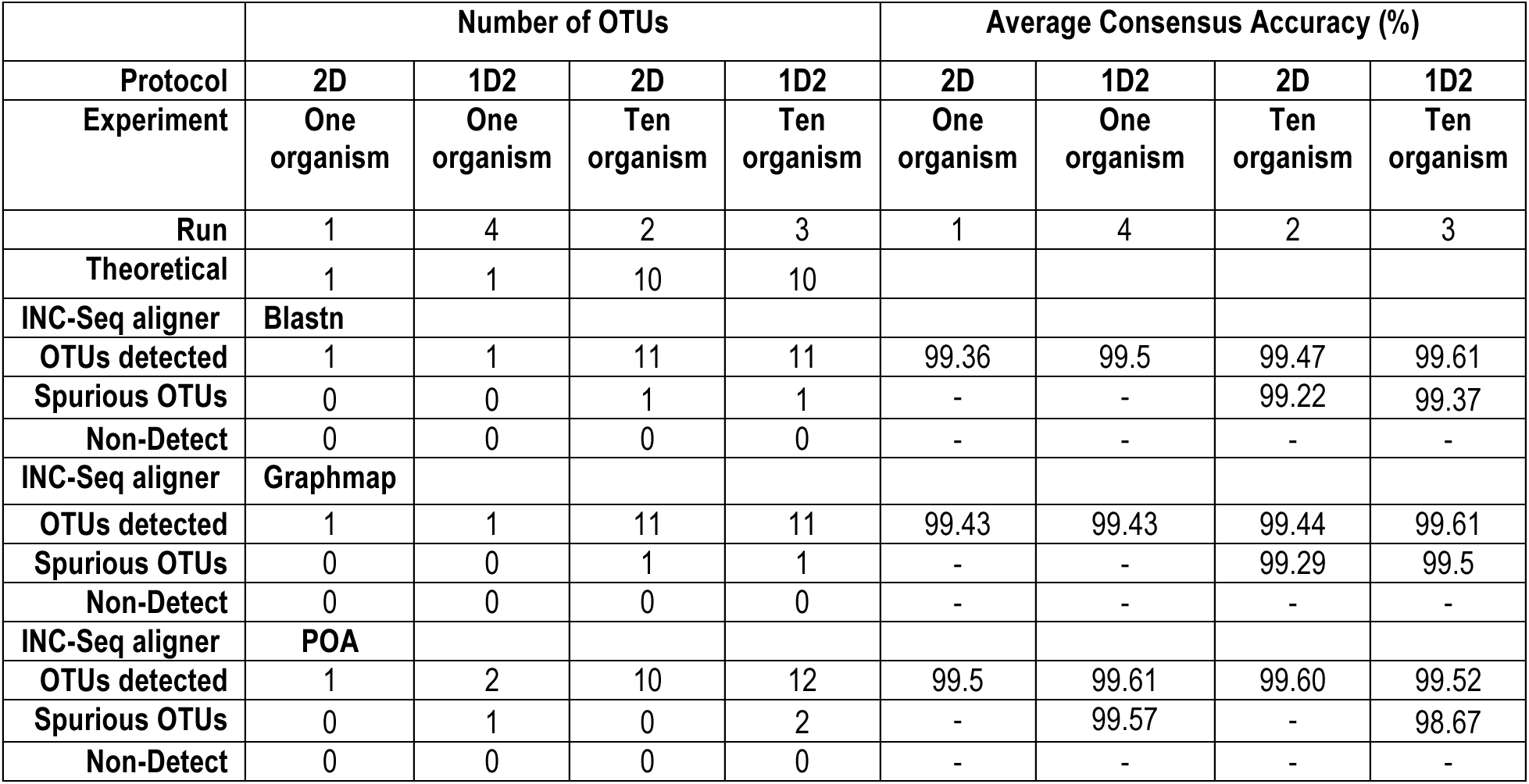
Number of OTUs detected and consensus sequence accuracy for all experiments using the nanoClust for OTU clustering and consensus calling approach.

The nanoClust approach with three partitions was far superior to the direct clustering of full-length reads and resulted in an accurate determination of the number of OTUs, with one-two spurious OTUs (when using 3 concatemers threshold for INC-Seq) and no false negatives (Table 2) depending on the type of experiment and INC-Seq aligner used. MAFFT-G-INS-i alignment of 50 reads from each OTU resulted in consensus reads with the entire read aligned to the reference, and average consensus read of 99.5% and accuracy values for individual OTUs ranging from 99.2 to 100%. Nearly all errors in the nanoClust consensus reads originate from single base pair deletions in a few homopolymers regions (homopolymers > 4 bp) and no detectable insertions, with one-two mismatches associated with the spurious OTUs (Figure 8). While accurate OTU estimation allowed for single OTUs to be detected in the one organism experiment, the overall community structure deviated from the theoretical community structure for the ten organism experiments (Figure S4) and thus additional protocol optimization is essential to ensure levels of deviation from theoretical community structure does not exceed what may be seen from PCR biases [33]. Phylogenetic analyses of the consensus sequences demonstrated close placement of the OTU consensus sequences with their corresponding references, with excellent pairwise alignment between the two (Supplementary Figure S3).

**Figure 8:**
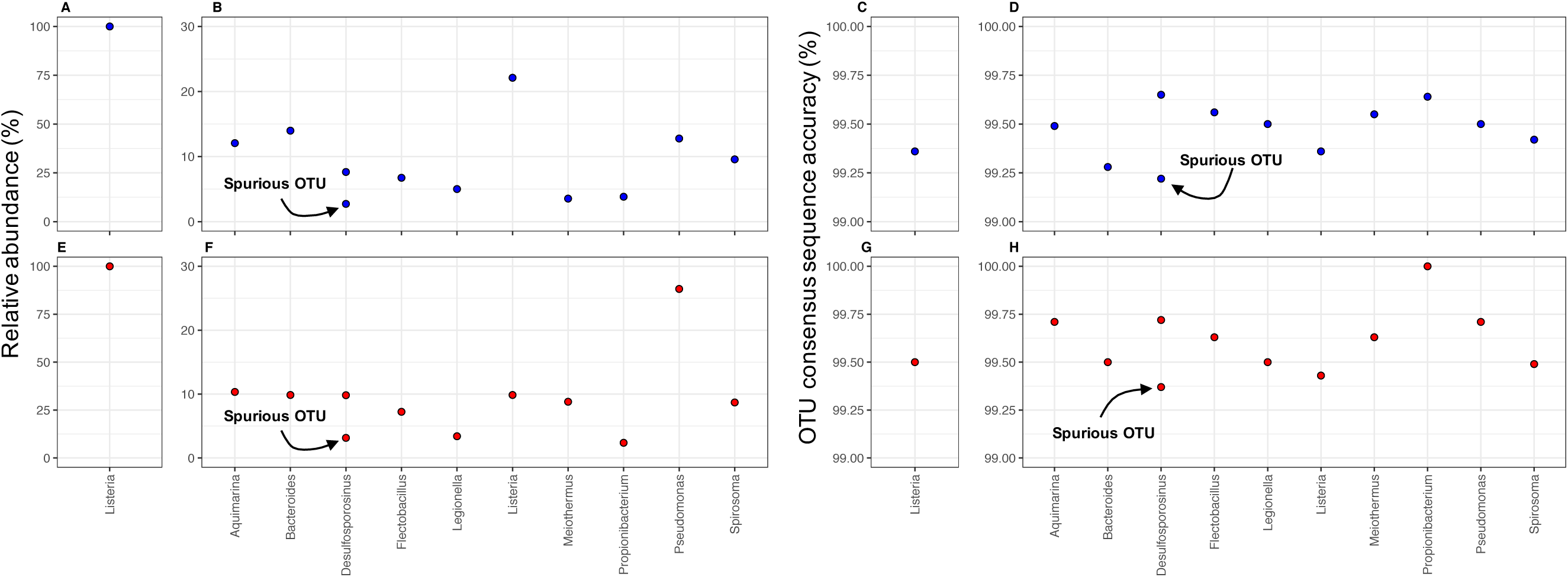
Relative abundance of OTUs for one (A, E) and ten organism experiments (B, F) for 2D (blue data points) and 1D2 (red data points) experiments post nanoClust when using blastn algorithm during INC-Seq. nanoClust clustering and consensus sequence generation resulted in few spurious OTUs and average similarity to the reference sequence of ~ 99.5%. Results are shown for one (C, G) and ten organism experiments (D, H) for 2D (blue data points) and 1D2 (red data points) experiments with the use of blastn during INC-Seq. The results were similar for Graphmap and POA alignment methods used during INC-Seq.

The nanoClust implementation in this study included a specified threshold of a maximum of 50 reads per OTU to generate OTU consensus sequences. This was feasible because our study focusses on a single organism and even mock community of ten organisms. Thus, the process of consensus construction was not limited by the number of reads that could be recruited. However, it would be critical to determine the potential for poor quality consensus sequence due to fewer reads with an OTU in naturally derived mixed microbial communities. To this end, we varied the number of reads used for consensus sequence construction from 5 to 100 for 2D and 1D2 data from the one organism experiment (Figure 9).

**Figure 9:**
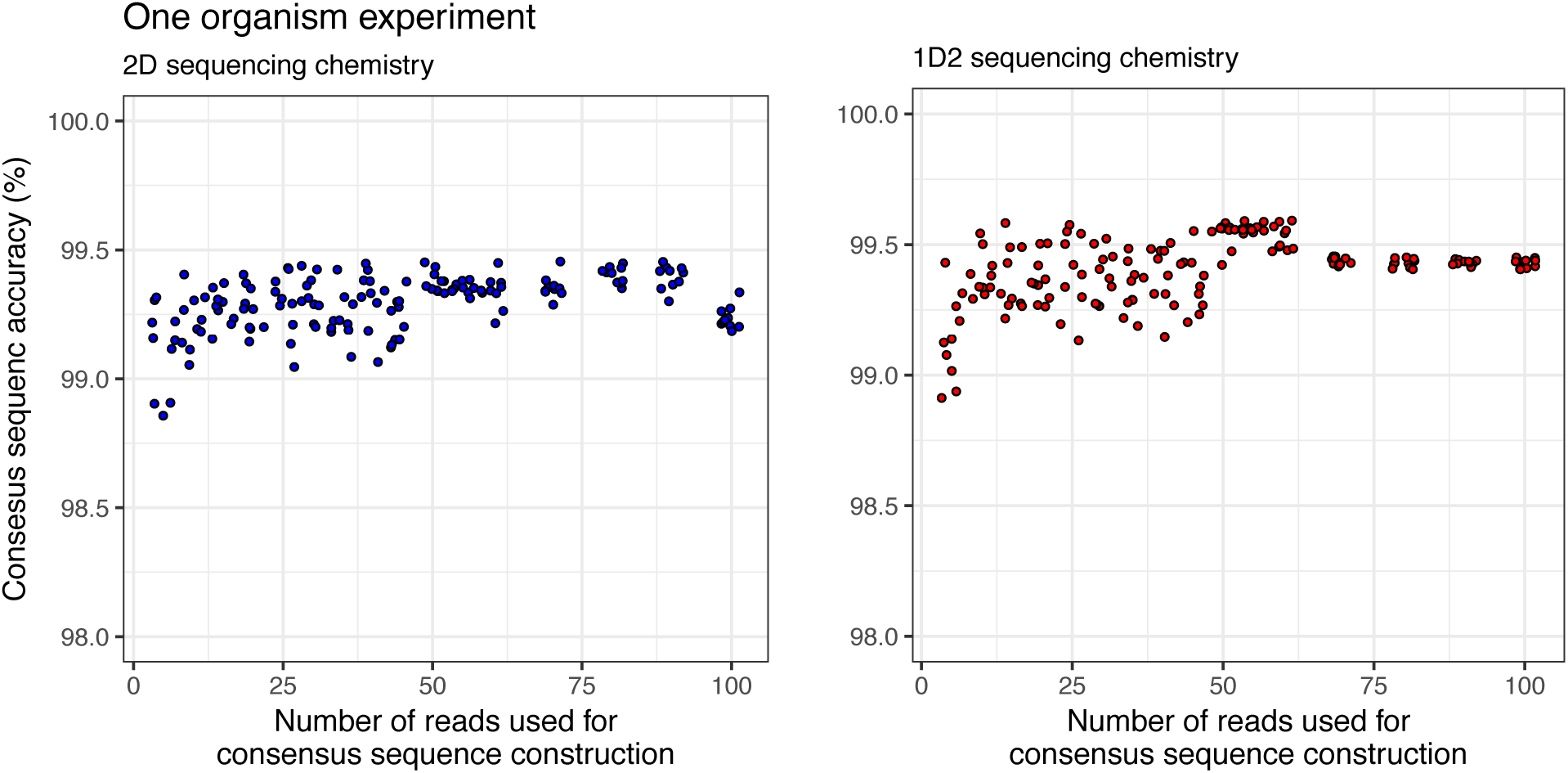
Consensus sequence accuracy plateaus with the use of 10–15 reads for MAFFT-G-INS-i alignment and consensus calling. However, with increasing number of reads used for consensus calling the variability in consensus sequence accuracy from repeated sampling of data diminishes significantly for both 2D and 1D2 sequencing chemistry. Data is shown for one organism experiment where the blastn aligner was used during INC-Seq.

The consensus sequence accuracy surpasses 99% with the use of more than five reads for consensus sequence construction and plateaus at approximately 10–15 reads. However, the variability in accuracy with repeated random sampling of data was much more pronounced when fewer than 50 reads were used for both 2D and 1D2 data. This suggests that consensus sequence accuracy is reliably high only for OTUs where a minimum of 50 reads are available for use in constructing the consensus sequence. This would have an impact on sequence quality of low abundance OTUs.

### NanoAmpli-Seq based improvements in sequence accuracy are not primarily associated with changes in nanopore sequencing chemistry

The INC-Seq study [21] utilized data generated from flow cells with R7 pores, while the present study used R9.4 and 9.5 pores with reported higher sequencing accuracy. Improvements in sequencing chemistry and basecalling allowed us to reduce the concatemer threshold for INC-Seq from six to three, which significantly increased the amount of data used for analysis. The second significant improvement of the updated sequencing chemistry is the much higher sequencing output. However, neither of these improvements result in improved data quality post INC-Seq processing alone (Figure 5). Thus, the chopSeq and nanoClust algorithms are critical for obtaining 99.5% sequence accuracy.

To demonstrate this, we re-processed the “Ladder replicate” data made available through the original INC-Seq study [21] using the NanoAmpli-Seq workflow. While re-analyzing R7 chemistry generated data from Li et al [21], we detected tandem repeats and incorrect primer orientation issue highlighted in this study. chopSeq was successfully able to remove tandem repeats and re-orient reads such that nearly the entire length of the read was now correctly aligned to the reference sequence for most of the reads. Thus, while INC-Seq reaches a median sequence accuracy of 97–98% (as described in the Li et al. [21]), post-processing by chopSeq improves read quality through read re-orientation and tandem repeat removal (Figure 10).

**Figure 10:**
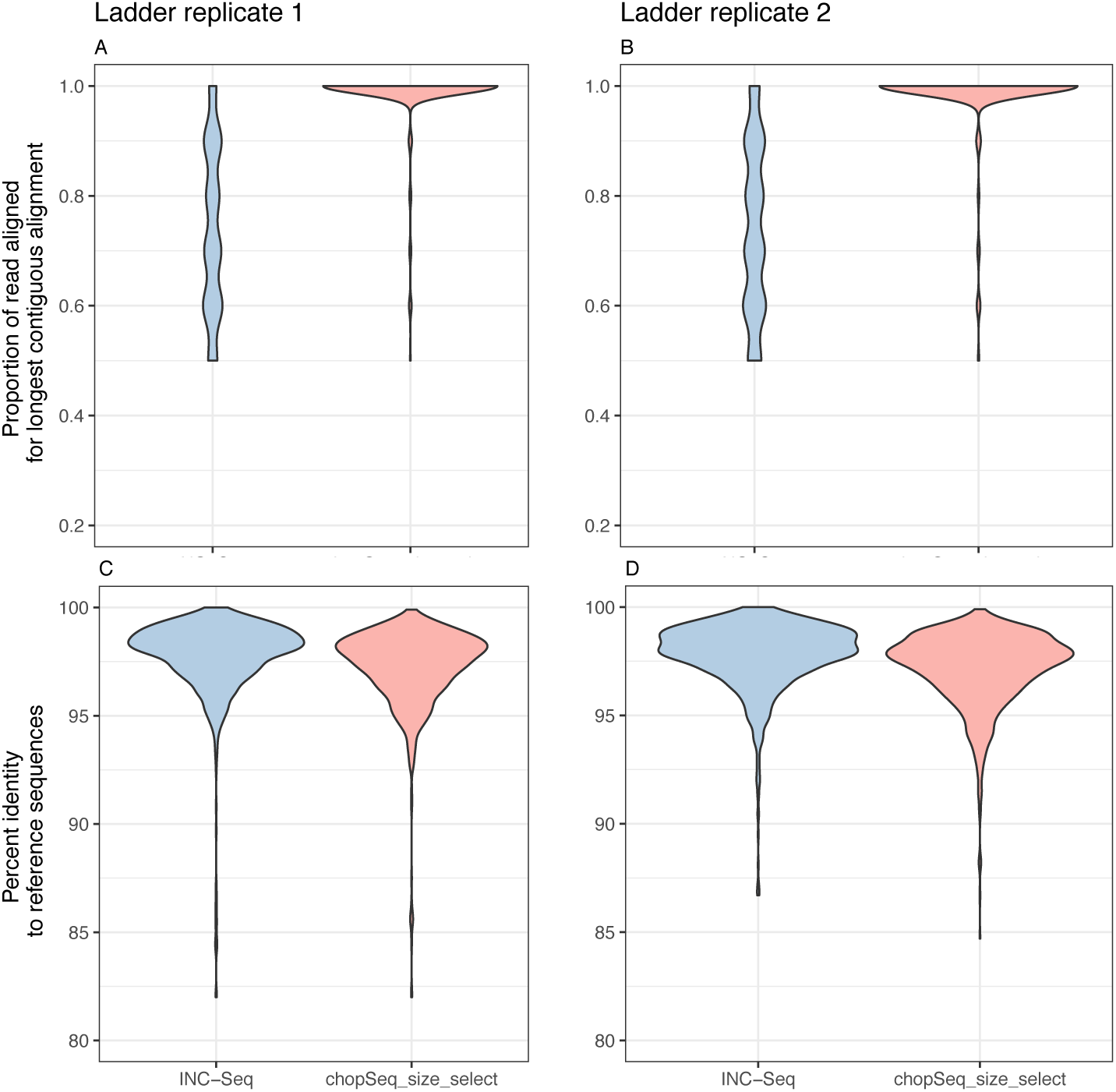
chopSeq based read re-orientation and tandem repeat removal allowed for nearly the entire length of the read to be aligned to the reference sequences (A, B) while maintaining the median sequence accuracy of INC-Seq consensus reads to ~97–98%.

Furthermore, nanoClust based clustering and consensus calling results in an average sequence accuracy of 99.5% for the data generated by Li et al [21]. The data generated by Li et al. [21] included the V3-V6 region of the 16S rRNA gene from 10-organism mock community with a staggered community structure including closely related organisms. This resulted in a theoretical number of 8 OTUs at 97% sequence similarity. Specifically, *Staphylococcus aureus* and *Staphylococcus epidermis* clustered into a single OTU at 97% sequence similarity (their 16S rRNA gene V3-V6 hypervariable regions are 98.8% similar to each other) and *Klebsiella pneumoniae* and *Salmonella typhimurium* clustered into a single OTU at 97% sequence similarity (their 16S rRNA gene V3-V6 hypervariable regions are 97.6 % similar to each other). Further, the Li et al. [21] only generated 2100 INC-Seq consensus reads combined for the two replicate sequencing runs. As a result, two of the low abundance OTUs with a relative abundance of 0.2% (*Neisseria*), and 0.1% (*Faecalibacterium*) were not detected after processing with chopSeq and nanoClust. This non-detection of low abundance OTUs is primarily a function of low sequencing depth rather than of the NanoAmpli-Seq workflow. Further, the sequence accuracy of most of the detected OTUs was in excess of 99.5%, while that of a single OTU (i.e., *Fusobacterium*) was 98.75% (Figure 11). Thus, we conclude that incorporating chopSeq correction of INC-Seq consensus reads followed by nanoClust based clustering and consensus calling was vital for improved sequence accuracy, irrespective of the changes in the sequencing chemistry.

**Figure 11:**
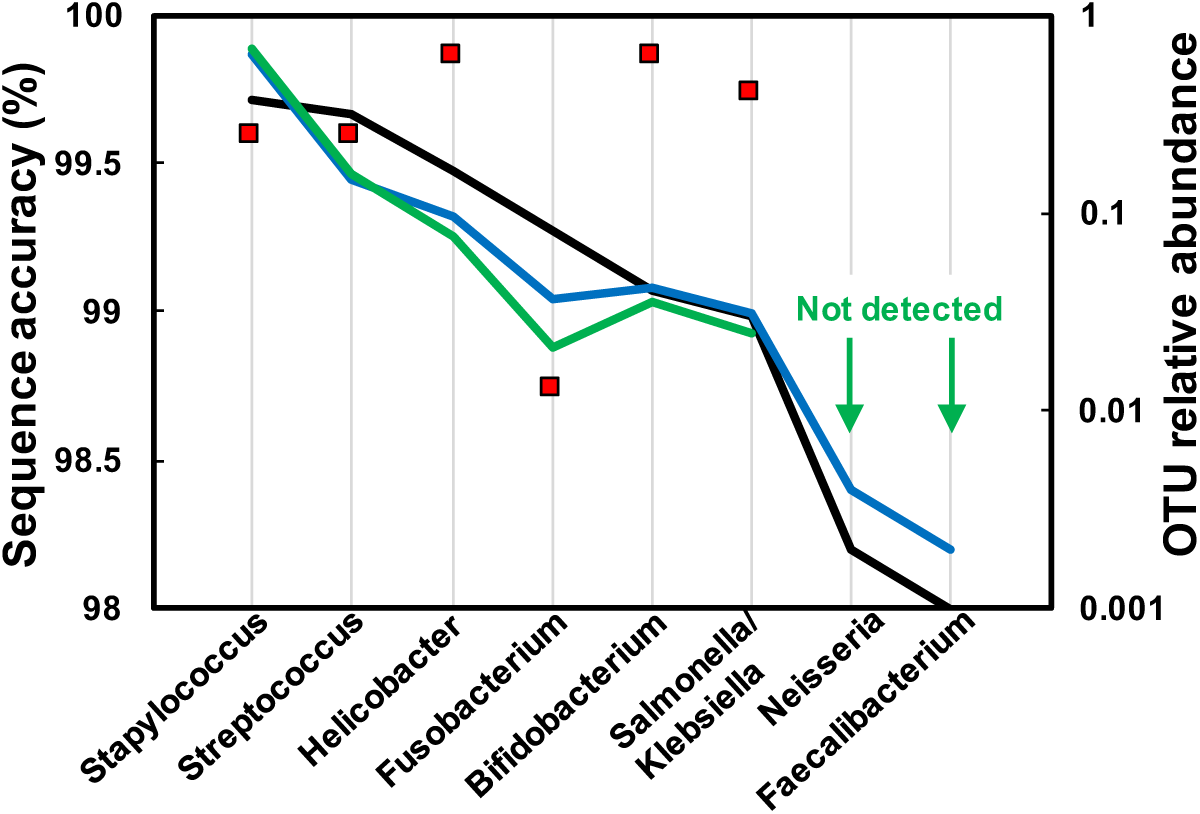
Sequence accuracy of detected OTUs post nanoClust for combined ladder replicate was ~99.5% (red squares). Two OTUs were not detected post nanoClust processing (green line) due to their low abundance (<0.5%) in the constructed mock community (black line). The blue line shows the relative abundance as reported by Li et al.

## Discussion and conclusions

The current study uses mock communities to develop and validate the NanoAmpli-Seq workflow for long amplicon sequencing on the nanopore sequencing platform. While this study focusses on the near full-length 16S rRNA gene, in principle the approach outlined by the NanoAmpli-Seq workflow should be amenable to amplicons generated from PCR amplification of any target gene irrespective of target gene length. While leveraging the previously described INC-Seq protocol, NanoAmpli-Seq adds several novel components which significantly enhances the amplicon sequencing workflow for the nanopore platform. The improvements over the previously described INC-Seq protocol involve modifications to both library preparation (i.e. PrimPol based primer synthesis for RCA, debranching and fragmentation, shorter protocol length) and the data analyses. Specifically, we identify and fix the issues associated with incorrect read-orientation and presence of tandem repeats in INC-Seq consensus reads, thus allowing for nearly the entire length of the chopSeq corrected reads to be aligned to the reference with accuracies (97–98%) similar to those described by Li et al [21]. While the original INC-Seq protocol prescribed a concatemer threshold of six, we halved the concatemer threshold to three. Thus, more than doubling the number of INC-Seq consensus reads available as a proportion of the base called reads. We could use a lower INC-Seq concatemer threshold due to both enhances in basecalling and sequencing accuracy of the nanopore platform [34]and the ability to perform another round of alignment and consensus calling during the nanoClust step.

The correction of INC-Seq consensus reads using chopSeq did not allow for sequences with high enough quality for direct OTU clustering using VSEARCH. However, the read partitioning-based sequence clustering allowed for accurate determination of the number of OTUs in the mock community. Further, *de novo* sequence clustering using nanoClust provided the opportunity to significantly increase the number of sequences used for consensus calling. In this study, we used 50 reads for consensus calling (i.e. 150x coverage considering three concatemer threshold set for INC-Seq) which resulted in average sequence accuracy of 99.5%. The use of more than 50 reads for consensus calling in nanoClust did not improve sequence accuracy while reducing the number of reads resulted in reduced precision. This threshold of 50 reads for both the 2D and 1D2 sequencing data suggests that the OTUs with fewer than 50 reads are likely to have sequence quality lower than those OTUs with greater than 50 reads. It should however be noted that using a 10 read threshold (i.e., 30x coverage when including three concatemer threshold for INC-Seq) consistently allowed for sequence accuracy consistently higher than 99% and thus sequence classification to species level using any of the current sequence classification approaches [35, 36] (i.e., RDP classifier) would be reliable even for lower abundance OTUs.

While the NanoAmpli-Seq workflow represents a significant improvement in amplicon sequencing on the nanopore platform, some fundamental limitations remain. For instance, the NanoAmpli-Seq sequence accuracy is still lower than those reported for short amplicons [3] or those generated from the assembly of SSU rRNA from metagenomic sequencing on the Illumina Platform [7, 9], and the full-length 16S rRNA sequencing on the PacBio platform using the approach described previously [17]. Our analysis shows that the sequence accuracy does not improve with more than 50 sequences used in the nanoClust based consensus calling process. Nearly all the errors in the OTU consensus sequences originate from single deletions at homopolymers regions, specifically for homopolymers greater than 4 bp. This homopolymer error issue on the nanopore platform is well known [14, 37] and is likely to best resolved during the base calling process rather than subsequent data processing or by processing signal data (rather than base called data) all the way through clustering, followed by base calling as the final step. The second limitation of our approach is the low data yield at the base calling step, i.e. base called reads represents only a small portion (i.e., 7–9% for 1D2 data) of the raw records. This data loss is significant and could potentially deter the widespread use of the nanopore platform for amplicon sequencing. While the precise cause of the low yield of pass reads post base calling is unclear, the proportion of pass reads in our study are not significantly different from those reported elsewhere. One current option would be to directly work with 1D rather than 1D2 data. However, the maximum sequence accuracy of 1D reads post INC-Seq consensus construction was only 94% and unsuitable for processing with chopSeq and nanoClust. The NanoAmpli-Seq workflow includes a *de novo* clustering step, and as long as the sequence accuracy post chopSeq is ~97% (3–10 concatemers required), the binning process should provide for sufficient coverage for consensus-based sequence correction to accuracies in excess of 99%. The final limitation of our approach is that the nanoClust relies on generating consensus sequences from multiple DNA sequences and thus there is the likelihood of clustering and generating a multi-species consensus from closely related species, i.e. those within 97% sequence similarity to each other. While we did not find evidence for this “multispecies consensus sequence” while analyzing data from Li et al. [21] which included closely related organisms, this possibility cannot be ignored. And thus, we recommend that researchers refrain from depositing NanoAmpli-Seq processed sequences in publicly available references databases, but utilize this approach for rapid screening of mixed microbial communities and limit the use of NanoAmpli-Seq processed data for within-study sample microbial community comparisons. Future improvement to avoid the likelihood of “multispecies consensus sequence” would be to utilize primers with barcodes consisting of random N bases (i.e., unique molecular tags), similar to that used by Karst et al. This could allow clustering of reads originating from the same original sequence using the unique molecular tags.

## Methods

### Mock community description and preparation

Two different mock communities were constructed for the experiments outlined in this study. First, a single organism mock community was constructed by amplifying the near full-length of the 16S rRNA gene from genomic DNA of *Listeria monocytogens* using primers sets 8F (5’- AGRGTTTGATCMTGGCTCAG-3’) and 1387 R (5’-GGGCGGWGTGTACAAG-3’), both with 5’ phosphorylated primers (Eurofins Genomics) ^26^. Phosphorylated ends are essential for the subsequent self-ligation step. PCR reaction mix was prepared in 25µl volumes with use of 12.5μl of Q5^®^ High-Fidelity 2X Master Mix (New England BioLabs Inc., M0492L), 0.8μl of 10pmol of each primer, 9.9μl of nuclease-free water, (Roche Ltd.) and 1ng of bacterial DNA in total followed by PCR amplification as described previously^26^. PCR amplicons from replicate PCR reactions were combined and purified with use of HighPrep™ PCR magnetic beads (MagBio, AC-60050) at 0.45x ratio. The ten organism mock community was constructed from purified near full-length 16S rRNA amplicons of 10 organisms. Briefly, genomic DNA from 10 bacteria were obtained from DSMZ, Germany (Supplementary Table S1) and the aforementioned primers, PCR reaction mix and thermocycling conditions were used to independently PCR amplify the near full-length 16S rRNA gene, followed by purification using HighPrep™ PCR magnetic beads as detailed above. The purified amplicons from each organism were quantified on the Qubit using dsDNA HS kit, normalized to 4ng/μl, and combined to generate an amplicon pool consisting of an equimolar proportion of the 16S rRNA gene amplicons of the 10 organisms.

### DNA sequencing library preparation

To circularize the linear amplicons into plasmid-like structures, 5μl of Blunt/TA Ligase Master Mix (New England Biolabs, M0367L) was added to 55 μl of amplicon pool at a concentration of 1ng/μl and incubated for 10min at 15°C then 10min at room temperature to (total time = 20 minutes). Not all linear amplicons self-ligate into plasmid-like structures, but some are likely to cause long chimeric linear amplicons. These long chimeric structures were removed using magnetic bead-based purification with the following modifications. HighPrep™ PCR magnetic beads were homogenized by vortexing followed by aliquoting 50µl into a sterile 2ml tube and placed on a magnetic rack for 3min. A total of 25μl of the supernatant was carefully removed using a sterile pipette to concentrate the beads to 2x its original concentration. The tube was removed from the magnetic rack and vortexed vigorously to resuspend the beads. This concentrated bead solution was used at a ratio of 0.35x to remove any amplicons greater than 2000 bp in the post-ligation reaction mix. Briefly, the post-ligation product was mixed with concentrated bead solution at the 0.35x ratio by vortexing followed by incubation for three minutes at room temperature. The tube was placed on the magnetic rack to separate the beads from solution, followed by transferring of clear liquid containing DNA structures less than 2000 bp into new sterile tubes. Sample containing short self-ligated molecules was subject to another round of concentration using standard magnetic beads at 0.5x ratios according to manufacturer instructions and eluted in 15μl of warm nuclease-free water. Concentrated and cleaned DNA pool consisting of plasmid-like structures and remaining linear amplicons was then processed with Plasmid-Safe™ ATP-Dependent DNase (Epicentre, E3101 K) reagents to digest linear amplicons using the mini-prep protocol according to manufacturer instructions and was followed by another round of cleanup with magnetic beads at 0.45x ratio as described before and then eluted in 15μl of warm nuclease-free water.

The pool containing plasmid-like structures was subject to RCA with use of TruPrime^TM^ RCA Kit (Sygnis, 390100) random hexamer-free protocol. Samples were prepared in triplicate and processed according to manufacturer protocol with all incubations performed in triplicate for 120–150 min depending on the assay efficiency. The progress of RCA was monitored by measuring the concentration of DNA using Qubit^®^ 2.0 Fluorometer at 90, 120 or 150 min time points. The negative control sample, consisting of reagents without any circularized plasmid-like amplicons, were processed and analyzed concomitantly with the samples. The final concentration of the RCA product after 150 minutes of incubation was typically 70 ng/μl when using a starting DNA concentration of 0.35–0.4 ng/μl with no detectable unspecific product formation in the negative control.

Replicates RCA products were combined (~4.5μg of DNA in total) and subject to de-branching and fragmentation of post-RCA molecules to remove hyperbranching structures generated during RCA. The RCA product was first treated with T7 endonuclease I enzyme (New England BioLabs, M0302S) by adding 2μl of the reagent to the 65μl of RCA product followed by vortexing and incubation as recommended by the manufacturer. Subsequently, the reaction mix was transferred into a g-TUBE (Covaris, 520079) and centrifuged at 1800 rpm for 4min or until the entire reaction mix passed through the fragmentation hole. The g-TUBE was reversed, and centrifugation process was repeated. Post debranching and fragmentation, short fragments were removed using the modified bead-based cleanup step using concentrated bead solution (see above for concentration procedure). Concentrated beads were mixed with fragmented RCA product at a 0.35x ratio, vortexed for 15sec, and incubated at room temperature for 3min then placed on a magnetic rack until the beads separated and the supernatant was removed. The beads were subsequently washed with 70% freshly prepared ethanol according to manufacturer protocols. Size-selected amplicons bound to the beads were eluted in 41μl of warm nuclease-free water. Preliminary experiments indicated that one round of de-branching did not completely resolve the hyperbranching structure, which was inferred based on poor sequencing yield likely caused due to pore blocking by hyperbranched DNA. As a result, a second round of enzymatic de-branching using T7 endonuclease I was added and the de-branched product was cleaned a second time using the bead-based clean-up step. Figure S4 shows an example BioAnalyzer traces of the RCA product post-debranching/fragmentation and post-cleanup using magnetic bead-based protocol.

Finally, the de-branched RCA product was treated with NEBNext^®^ FFPE DNA Repair Mix (New England BioLabs, M6630S) for gap filling and DNA damage repair caused during g-TUBE fragmentation and T7 endonuclease I enzyme. All reagent components were combined with de- branched RCA product according to manufacturer recommendations and incubated at 12°C for 10min then at 20°C for another 10min. Post incubation, the repaired RCA product was cleaned using standard magnetic beads at 0.5x ratio, washed with 70% ethanol, and eluted in 46μl of warm nuclease-free water. The concentration of the DNA product was measured using Qubit and was approximately 20–25 ng/μl with a total yield of ~1000 ng of DNA with product size typically ranging from 1500bp to 20,000 bp. A total of 45μl DNA pool of concatamerized amplicons was prepared for sequencing using the standard 2D and 1D2 library preparation protocol by Oxford Nanopore Technologies (SQK-LSK208, SQK-LSK308) according to manufacturer specifications to obtain pre-sequencing mix. Moreover, the final concentration for prepared libraries was determined using dsDNA HS kit on the Qubit instrument. A detailed step-by-step protocol is provided in the supplementary text.

### DNA sequencing

The MinION MkIB was connected to Windows personal computer compatible with Oxford Nanopore Technologies requirements. R9.4 (FLO-MIN106) and R9.5 (FLO-MIN107) flow cells were placed onto the MinION Mk1B (Oxford Nanopore Technologies). Platform quality control was performed using MinKNOW software (v1.4.2 for 2D and v1.6.11 for 1D2 libraries). Only flow cells containing above 1100 active pores were used in this study. Each flow cell was primed twice according to ONT specifications using priming buffer consisting of equal parts of running buffer (RBF1) and nuclease-free water with 10 min breaks between subsequent primes. The loading mix was prepared with a 12μL pre-sequencing mix, 75μL of RBF1, and 63μL of nuclease-free water. Loading mix was sequenced with MinKNOW settings appropriate for 2D or 1D2 options and standard 48h processing time for every run. Albacore 1.2.4 was used to convert raw signals into HDF5 file format using switch options FLO-MIN106 and SQK-LSK208 for 2D data and FLO-MIN107 and SQK-LSK308 for 1D2 data.

### Data processing

HDF5 raw signals from each sequencing run were converted to FASTQ format using Fast5-to-Fastq (https://github.com/rrwick/Fast5-to-Fastq) and then from FASTQ to FASTA with seqtk (https://github.com/lh3/seqtk). The resultant data was subject to INC-Seq (https://github.com/CSB5/INC-Seq) processing using blastn, Graphmap, and POA aligners with concatemer threshold of 3 with the iterative flag for consensus error correction using PBDAGCON. INC-Seq consensus reads were subject to chopSeq by specifying forward and reverse primer sequences and upper (1450 bp) and lower (1300 bp) read size thresholds for size selection (chopSeq_SS). The chopSeq processed reads were subsequently processed using nanoClust with partition size limits (flag “-s”) of 0,450,451,900,901,1300 which splits the reads into three partitions of 450, 450, and 400 bp respectively prior to further processing. VSEARCH was used for chimera removal (using uchime) followed by clustering of reads in each partition at a hardcoded sequence similarity threshold of 97%. For the optimal binning results, nanoClust then outputs consensus sequence for each OTU based on MAFFT-G-INS-i pairwise alignment as described previously and outputs an OTU table with reads corresponding to each OTU after discarding singletons.

Fasta files from all stages (i.e. raw, INC-Seq, chopSeq, chopSeq_SS, and OTU consensus sequence) were analyzed for read lengths in R [38]. The number of concatemers on each raw read was estimated by dividing the length of each raw read with the length of its corresponding INC-Seq consensus read. At each stage of processing, the reads were aligned to reference dataset using blastn, and only the match with the highest bitscore was considered. The ratio of read to reference alignment at appropriate points (as discussed above) was estimated based on blastn results. The percent identity from the blastn results were used to measure consensus sequence accuracy for the nanoClust output. All figures for the manuscript were generated in R using packages “ggplot2”, “gridExtra”, and “cowplot” (https://github.com/wilkelab/cowplot), as appropriate. Neighbor Joining tree construction (Figure S3 and S4) was performed after muscle alignment [39] (default parameters) and using Jukes-Cantor model was constructed in Geneious (version 8) using 100 bootstraps.

## Availability and requirements

Project name: NanoAmpli-Seq

Project home page: https://github.com/umerijaz/nanopore

Operating system: Linux

Programming language: Python

## Availability of supporting data

All data is available on European Nucleotide Archive (ENA) under primary accession number: PRJEB21005.

## Abbreviations

ONT, Oxford Nanopore Technologies; RCA, Rolling circle amplification; SMRT, single molecular real-time; INC-Seq, Intramolecular-ligated Nanopore Consensus Sequencing

## Declarations

## Acknowledgements

This research was supported by EPSRC award no: EP/M016811/1. STC is supported by the EPSRC Doctoral Training Center at University of Glasgow. UZI is supported by a NERC Fellowship (NE/L011956/1).

## Author contribution

STC, UZI, and AJP designed and developed the experiments. STC performed the experiments. UZI wrote the code for chopSeq and nanoClust. STC, UZI, and AJP analyzed the data and contributed equally to writing this manuscript.

## NanoAmpli-Seq: A workflow for amplicon sequencing for mixed microbial communities on the nanopore sequencing platform

Szymon T Calus, Umer Z Ijaz, Ameet J Pinto

## Protocol for sample preparation for NanoAmpli-Seq workflow

**Consumables required**:

1. Sterile Filtered pipette tips from any vendor
2. 0.2 ml PCR grade tubes from any vendor
3. Wide Bore Filtered pipette tips from any vendor
4. Qubit dsDNA HS Assay kit. Vendor: ThermoFisher Scientific. Catalog number: Q32851.
5. Primers for PCR amplification of 16S rRNA gene with phosphorylated 5’ ends can be ordered from any provider:

a. Forward primer: 8F: [PHO] AGRGTTTGATCMTGGCTCAG
b. Reverse primer: 1387 R: [PHO] GGGCGGWGTGTACAAGRC
6. The master mix for PCR amplification: Q5^®^ High-Fidelity 2X Master Mix. New England Biolabs, Inc. Catalog number: M0492S
7. Nuclease-free Water. Vendor: New England Biolabs, Inc. Catalog number: B1500S
8. HighPrep^TM^ PCR paramagnetic bead solution. Vendor: MAGBIO. Catalog number: AC- 60050
9. 70% ethanol (prepared from denatured ethanol).
10. Magnetic stand for 1.5 ml tubes. MagStrip Magnet Stand 10. Vendor: MAGBIO. Catalog number: MBMS-10
11. Blunt/TA Ligase Master Mix. Vendor: New England Biolabs, Inc. Catalog number: M0367S
12. DNA LoBind Tubes, 1.5 ml. Vendor: Eppendorf. Catalog number: 003018078
13. Plasmid-Safe^TM^ ATP-Dependent DNAse. Vendor: Epicentre. Catalog number: E3101 K.
14. TruePrime^TM^ RCA kit. Vendor: Expedeon. Catalog number: 390100
15. T7 endonuclease I. Vendor: New England Biolabs, Inc. Catalog number: M0302S.
16. g-TUBE. Vendor: Covaris. Catalog number: 010145
17. NEBNext^®^ FFPE DNA Repair Mix. Vendor: New England Biolabs Inc. Catalog number: M6630S
18. Ligation Sequencing Kit 1D. Catalog number: SQK-LSK108 and/or Ligation Sequencing Kit 1D2. Catalog number: SQK-LSK308. Vendor: Oxford Nanopore Technologies
19. Flow Cell (R9.4) and/or Flow Cell (R9.5). Vendor: Oxford Nanopore Technologies

**Equipment required:**

1. PCR thermocycler from any vendor
2. PCR hood
3. Thermal mixer with appropriate blocks from any vendor
4. Pipettes from varying volumes range from any vendor
5. Centrifuge for 2 ml and 0.2 ml tubes from any vendor
6. MinION^TM^ Mk1b device and compatible personal computer

**Step 1: PCR amplification of 16S rRNA gene.**

1. Combine the following components using volumes below or as appropriate for your experiment in a PCR tube.

1. Q5^^®^^ High-Fidelity 2X Master Mix: 12.5 μl
2. Forward primer (10 pmol): 0.8 μl
3. Reverse primer: (10 pmol): 0.8 μl
4. Template DNA: 1 μl
5. Nuclease-free water: 9.9 μl.
2. Amplify PCR reaction mix at the following PCR conditions:

| Segment | Temp (°C) | Time (sec) | Cycles |
| --- | --- | --- | --- |
| Initial denaturation | 98 | 00:30 | 1 |
| Denaturation | 98 | 00:05 | 20 |
| Annealing | 59 | 00:10 |  |
| Extension | 72 | 00:35 |  |
| Final extension | 72 | 02:00 | 1 |
| Hold | 8 | 00:00 | 1 |

**Step 2: PCR product clean up.**

1. Follow vendor instructions (MAGBIO) to clean PCR product at 0.45x bead ratio as described at the following URL: http://www.magbiogenomics.com/image/data/Literature/Protocols/HighPrep%20PCR%20Protocol.pdf

**Step 3: PCR product concentration estimation.**

1. Follow vendor instructions (ThermoFisher) to determine the concentration of cleaned PCR product using Qubit^TM^ dsDNA HS Assay kit as described at the following URL: https://assets.thermofisher.com/TFS-Assets/LSG/manuals/Qubit_dsDNA_HS_Assay_UG.pdf

**Step 4: Self-ligation for the formation of plasmid-like structure.**

1. Dilute PCR product from step 2 to 2–3 ng/μl using nuclease-free water
2. Mix 55 μl of diluted PCR product with 5 μl of Blunt/TA Ligase Master Mix in 0.2 ml PCR grade tubes
3. Gently mix by flicking the tube a few times
4. Centrifuge tube for 10 seconds
5. Incubate for 10 min at 15°C in a PCR thermocycler
6. Gently mix by flicking the tube a few times
7. Centrifuge tube for 10 seconds
8. Incubate for another 10 min at room temperature.

**Step 5: Reverse phase cleanup.**

1. Vortex HighPrep^TM^ PCR paramagnetic bead solution
2. Transfer 50 μl of bead solution into clean DNA LoBind 1.5 ml tube
3. Place the tube on a magnetic rack until beads separate from the liquid
4. While the tube is on the magnetic rack, remove 25 μl of liquid from the tube using sterile pipette tip, taking care not to disturb the beads
5. Remove the tube from magnetic rack and gently vortex to resuspend the beads.
6. Add the self-ligation mix from step 4 to concentrated beads at a ratio of 0.35x bead ratio.
7. Incubate the mixture for 3 minutes at room temperature.
8. Place tube on a magnetic rack
9. Allow the beads to separate from the liquid. The beads contain long linear amplicons (potentially chimeric amplicons) while the liquid contains short linear amplicon and plasmid-like structures. Carefully remove the clear liquid from the tube and move it to step 6.

**Step 6: Plasmid and short amplicon clean-up.**

1. Follow vendor instructions (MAGBIO) to clean plasmid and short amplicon mix at 0.45x bead ratio as described at the following URL: http://www.magbiogenomics.com/image/data/Literature/Protocols/HighPrep%20PCR%20Protocol.pdf

**Step 7: Removal of linear molecules from plasmid mix.**

1. Combine 42 μl of clean product from step 6 (eluted in nuclease-free water) with 2 μl of 25 mM ATP solution, 5 μl of 10X reaction buffer and 1 μl of Plasmid-Safe DNASe in 0.2 ml PCR grade tube. The last three ingredients are provided with the Plasmid-Safe^TM^ ATP-Dependent DNAse kit.
2. Incubate at 37°C for 15 minutes in a PCR thermocycler
3. Clean product as described in Step 2
4. Determine the concentration of DNA in the cleaned product as described in Step 3.

**Step 8: Rolling Circle Amplification (RCA).**

Perform RCA in triplicate for each sample and include negative controls using nuclease-free water instead of cleaned product from step 7. Below are reaction conditions and volumes for a single RCA reaction:

1. Combine 2.5 μl of cleaned product from step 7 with 2.5 μl of Buffer D (provided with TruePrime^TM^ RCA kit) in 0.2 ml PCR grade tube and incubate at room temperature for 3–5 minutes
2. While the sample is being incubated, prepare the amplification mix consisting of 9.3 μl of nuclease-free water, 2.5 μl of reaction buffer, 2.5 μl of dNTPs, 2.5 μl of Enzyme 1 and 2.5 μl of Enzyme 2. All ingredients are included in the TruePrime^TM^ RCA kit
3. After 10 minutes, add 2.5 μl of Buffer N (provided with TruePrime^TM^ RCA kit) and prepared amplification mix to the tube
4. Mix by pipetting and incubate tube at 30°C for 150 minutes.
5. Follow Step 3 to determine the concentration of DNA in samples and negative controls.

**Step 9: Enzymatic de-branching.**

1. Combine triplicate reactions from step 8 into a single 0.2 ml PCR grade tube
2. Mix thoroughly to prepare single RCA product per sample
3. Combine 65 μl of RCA product with 2 μl of T7 endonuclease I and incubate at room temperature for 5 minutes.

**Step 10: Mechanical fragmentation.**

1. Transfer enzymatically de-branched RCA product into g-TUBE using wide bore pipette tips
2. Centrifuge at 1800 rpm for 4 minutes or until entire reaction mix passes through the fragmentation hole
3. Reverse the g-TUBE and centrifuge again at 1800 rpm for 4 minutes or until entire reaction mix passes through the fragmentation hole
4. Clean product with as described in Step 5, with the exception that the clear liquid after bead separation is discarded and bead-bound DNA is eluted according to vendor outlined protocol
5. Repeat step 9 (i.e. Enzymatic de-branching) on the g-TUBE fragmented product.
6. Clean product as described in Step 5, with the exception that the clear liquid after bead separation is discarded and bead-bound DNA is eluted according to vendor outlined protocol.

**Step 11: DNA Damage repair.**

1. Combine 53.5 μl of product from Step 10 with 6.5 μl of FFPE DNA Repair Buffer and 2 μl of NEBNext FFPE Repair mix in a 0.2 ml tube
2. Incubate at 20°C for 15 minutes
3. Clean product as described in Step 2
4. Quantify DNA concentration as described in Step 3.

**Step 12: DNA library preparation and nanopore sequencing.**

1. Prepare 2D or 1D2 libraries for nanopore sequencing according to the protocols described for the SQK-LSK108 or SQK-LSK308 kits by Oxford Nanopore Technologies.
2. Primer appropriate flow cell using protocols outlined by Oxford Nanopore Technologies.
3. Finally, sequence the prepare libraries on appropriate flow cells as outlined by Oxford Nanopore Technologies.

**Table S1:**
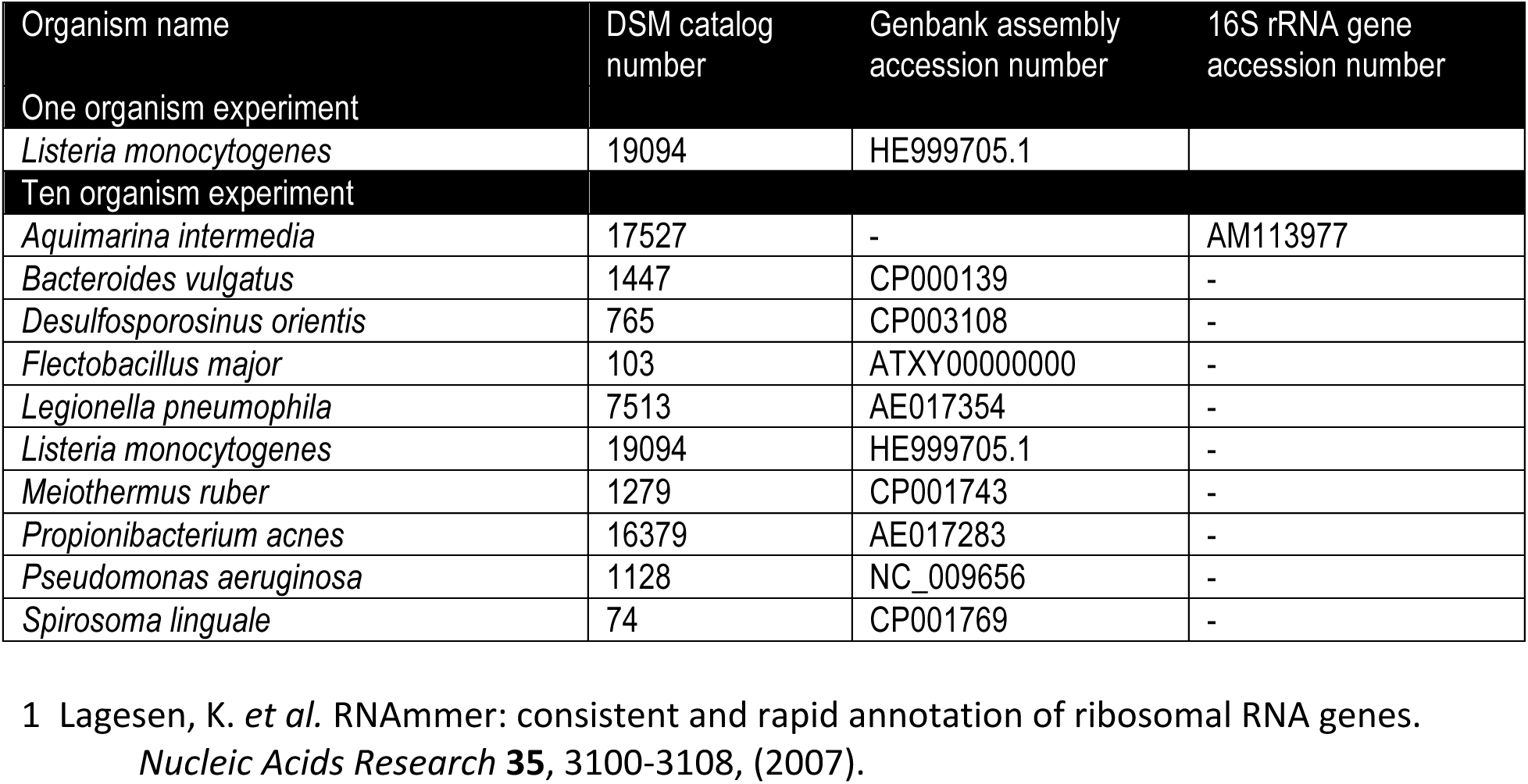
Names and DSM catalog numbers of bacteria used to construct mock communities for the single organism and ten organism experiments and their corresponding accession numbers are shown below. The 16S rRNA genes were extracted from genome assemblies using RNAmmer^1^ for bacteria for use in estimation of sequencing accuracy. Where genome assemblies were unavailable, the 16S rRNA gene sequence in Genbank was utilized.

**Figure S1:**
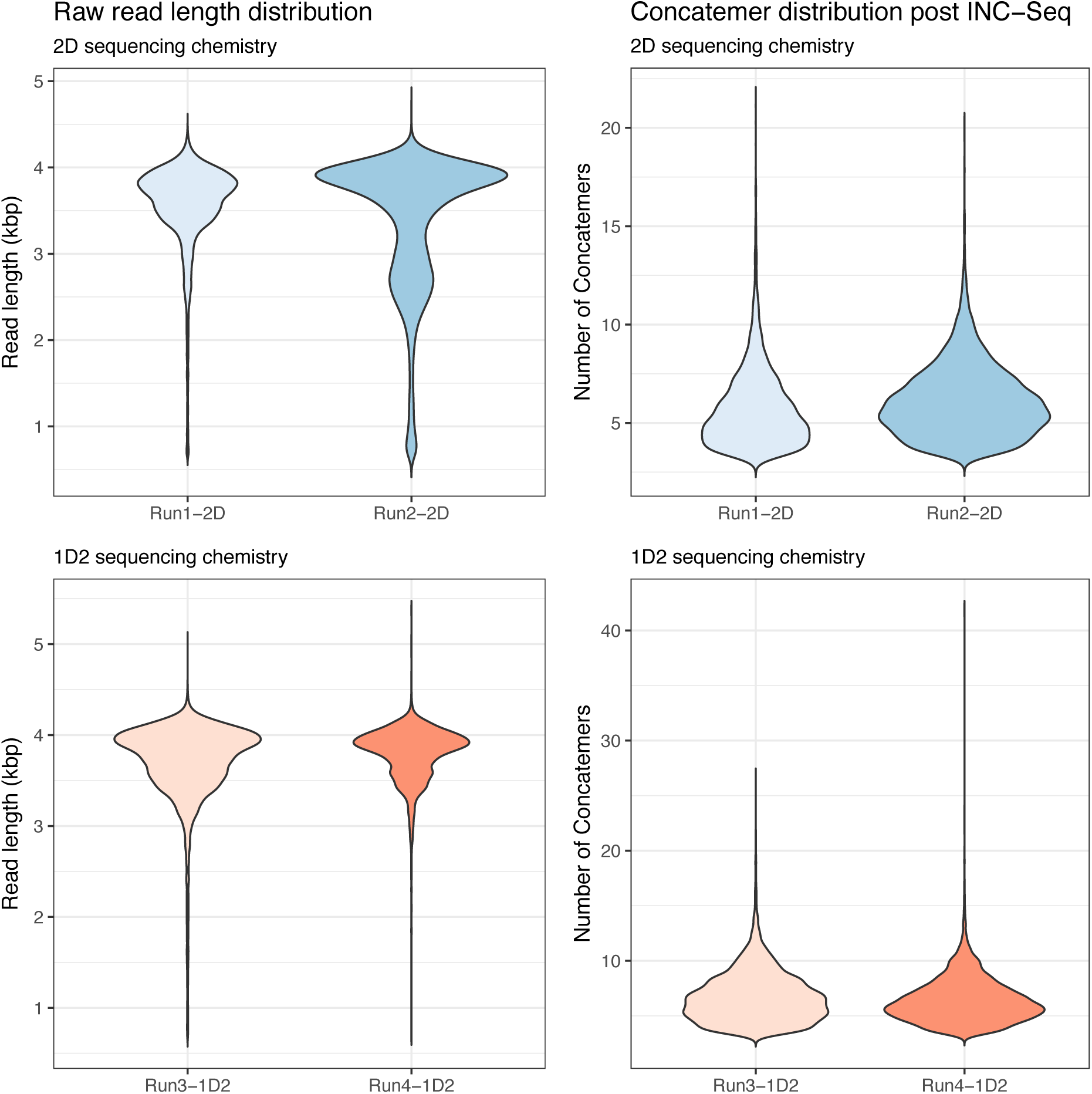
Violin plots showing the read length distribution of raw data (i.e., post base calling with Albacore 1.2.4), and the number of concatemers on each base called read estimated using read lengths from INC-Seq processing using the “blastn” alignment approach for all four experiments involving both 2D and 1D2 chemistry.

**Figure S2:**
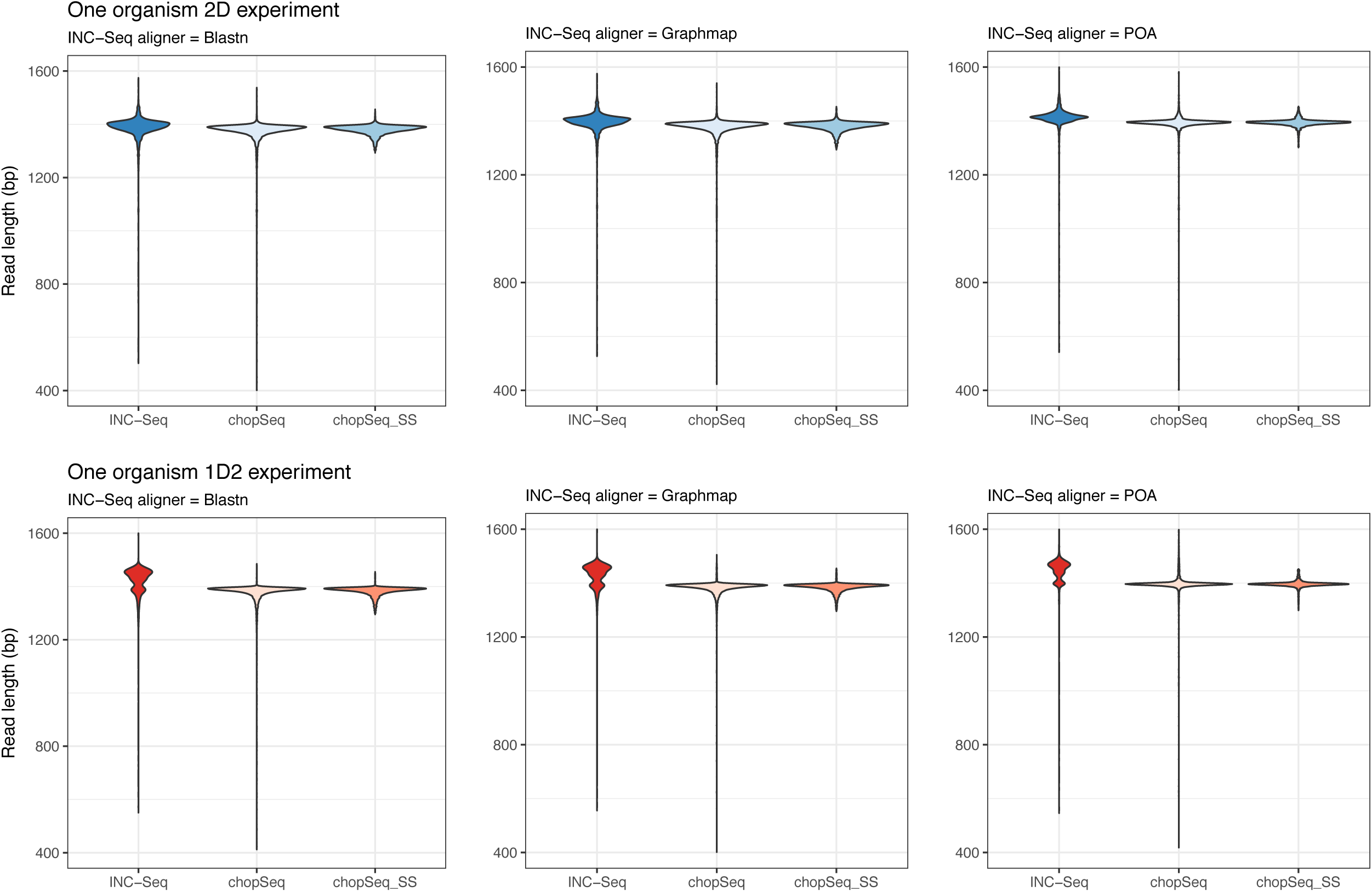
Violin plot of read length distributions for INC-Seq processed reads, chopSeq corrected reads after tandem repeat removal, and post-size-selection of chopSeq corrected reads and size selected reads (chopSeq_SS) for one organism experiments using both 2D and 1D2 sequencing chemistry.

**Figure S3:**
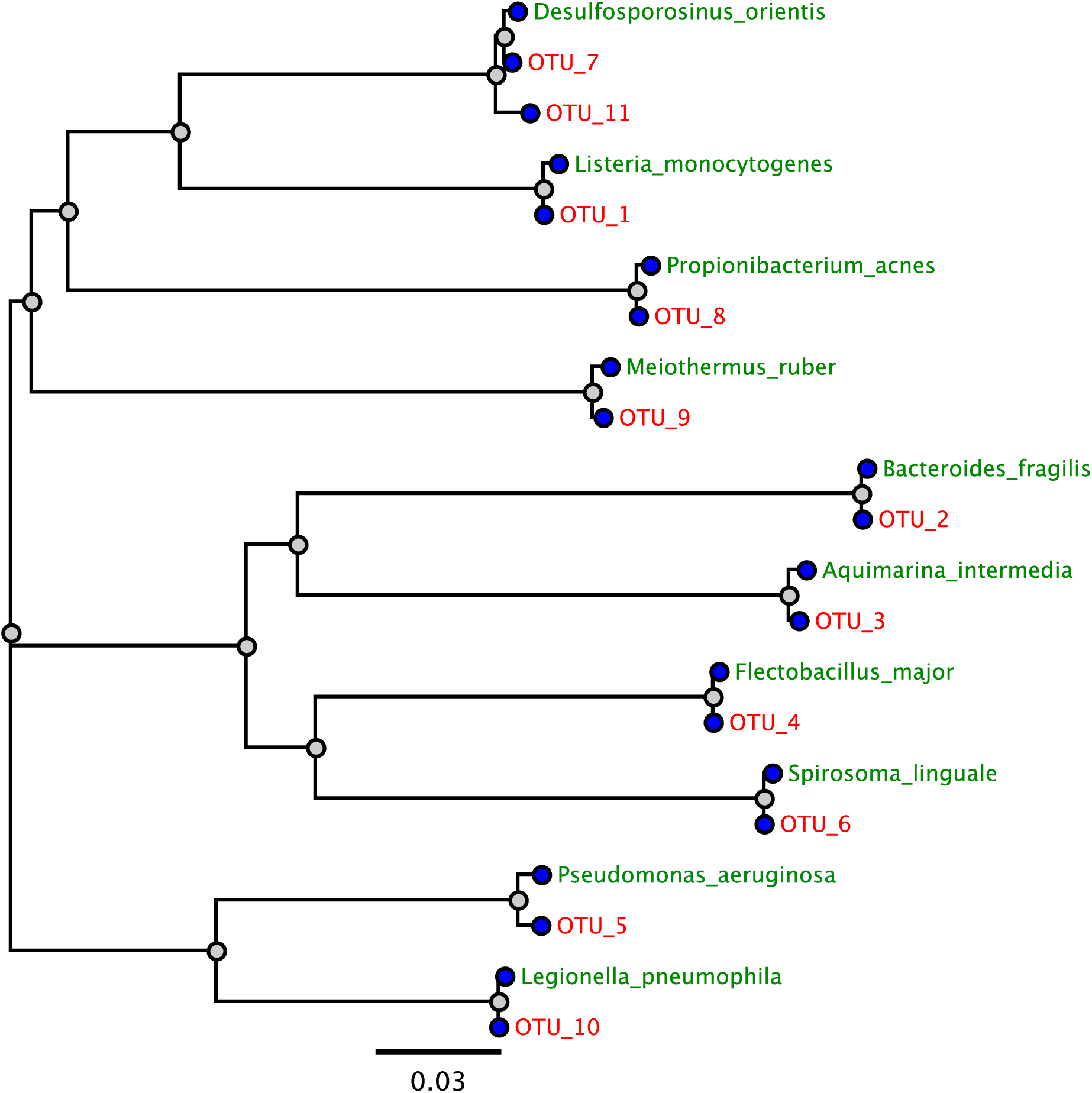
Consensus sequences from OTUs from Run 3 (1D2 sequencing chemistry, INC-Seq aligner: blastn) were combined with reference sequences and aligned using muscle (default parameters) and Neighbor-Joining tree using Jukes-Cantor model was constructed in Geneious (version 8) using 100 bootstraps. Reference sequences are labelled in green, and OTUs are labelled in red.

**Figure S4:**
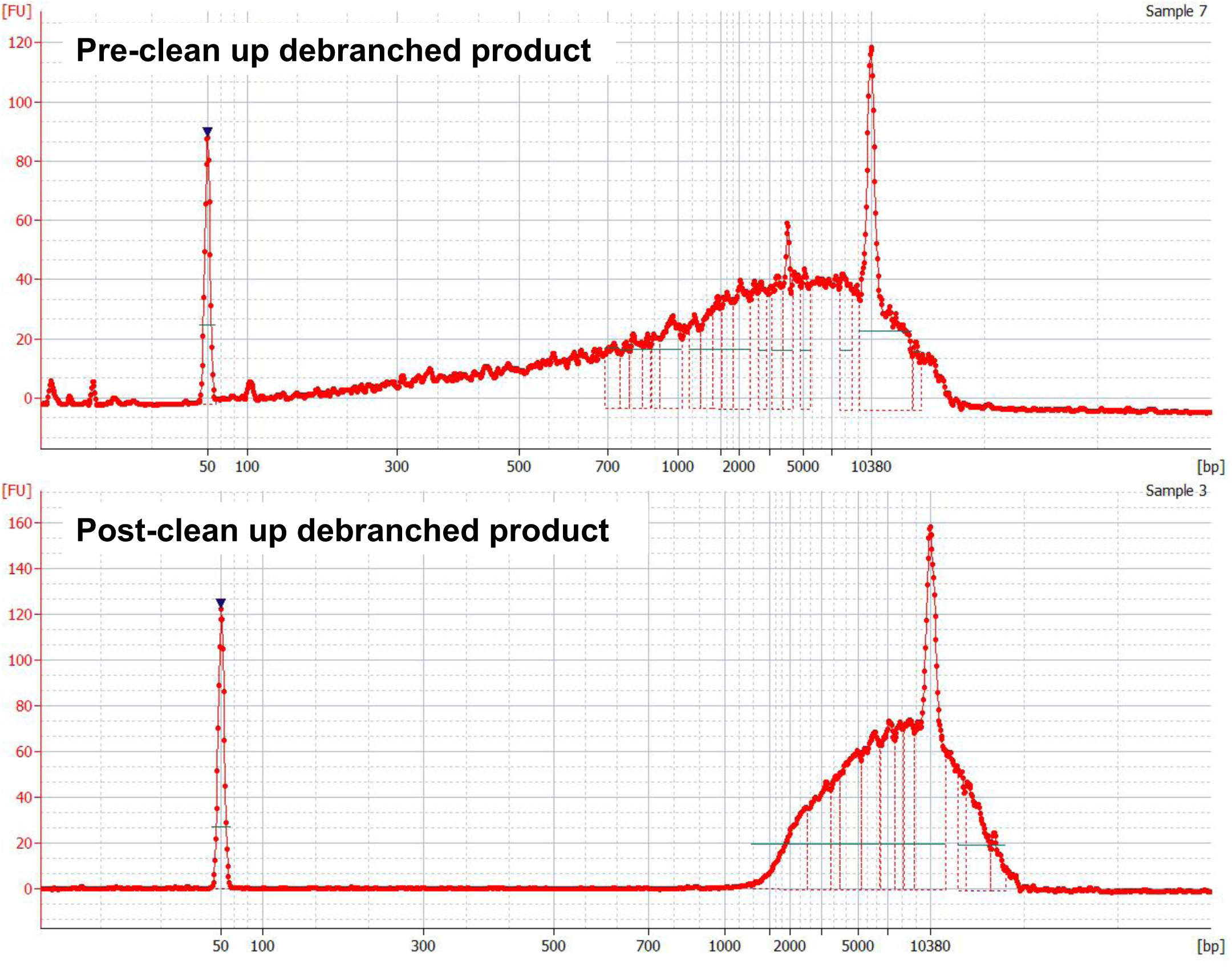
Example Bioanalyzer traces of de-branched product pre- and post-cleanup. The post-cleanup product was then treated for gap filling and DNA damage repair prior to being used for library preparation of sequencing according to ONT protocol for 2D and 1D2 chemistries.

## References

1. Caporaso JG, Lauber CL, Walters WA, Berg-Lyons D, Huntley J and Fierer N. Ultra-high-throughput microbial community analysis on the Illumina HiSeq and MiSeq platforms. ISME J. 2012;6 doi:10.1038/ismej.2012.8.

2. Caporaso JG, Lauber CL, Walters WA, Berg-Lyons D, Lozupone CA, Turnbaugh PJ, et al. Global patterns of 16S rRNA diversity at a depth of millions of sequences per sample. Proceedings of the National Academy of Sciences. 2011;108 Supplement 1:4516–22. doi:10.1073/pnas.1000080107.

3. Kozich JJ, Westcott SL, Baxter NT, Highlander SK and Schloss PD. Development of a dual-index sequencing strategy and curation pipeline for analyzing amplicon sequence data on the MiSeq Illumina sequencing platform. Appl Environ Microbiol. 2013;79 doi:10.1128/aem.01043-13.

4. Schoch CL, Seifert KA, Huhndorf S, Robert V, Spouge JL, Levesque CA, et al. Nuclear ribosomal internal transcribed spacer (ITS) region as a universal DNA barcode marker for Fungi. Proceedings of the National Academy of Sciences. 2012;109 16:6241–6. doi:10.1073/pnas.1117018109.

5. Goodwin S, McPherson JD and McCombie WR. Coming of age: ten years of next- generation sequencing technologies. Nat Rev Genet. 2016;17 6:333–51. doi:10.1038/nrg.2016.49.

6. Yuan C, Lei J, Cole J and Sun Y. Reconstructing 16S rRNA genes in metagenomic data. 1367–4811 (Electronic) doi:10.1093/bioinformatics/btv231.

7. Miller C, Baker B, Thomas B, Singer S and Banfield J. EMIRGE: reconstruction of full-length ribosomal genes from microbial community short read sequencing data. Genome Biology. 2011;12 5:R44.

8. Burke CA-O and Darling AE. A method for high precision sequencing of near full-length 16S rRNA genes on an Illumina MiSeq. 2167–8359 (Print).

9. Miller CS, Handley KM, Wrighton KC, Frischkorn KR, Thomas BC and Banfield JF. Short-Read Assembly of Full-Length 16S Amplicons Reveals Bacterial Diversity in Subsurface Sediments. PLOS ONE. 2013;8 2:e56018. doi:10.1371/journal.pone.0056018.

10. Karst SM, Dueholm MS, McIlroy SJ, Kirkegaard RH, Nielsen PH and Albertsen M. Retrieval of a million high-quality, full-length microbial 16S and 18S rRNA gene sequences without primer bias. Nature Biotechnology. 2018;36:190. doi:10.1038/nbt.4045.

11. Mardis ER. DNA sequencing technologies: 2006–2016. Nature Protocols. 2017;12:213. doi:10.1038/nprot.2016.182.

12. Wommack KE, Bhavsar J and Ravel J. Metagenomics: Read Length Matters. Applied and Environmental Microbiology. 2008;74 5:1453–63. doi:10.1128/aem.02181-07.

13. Koren S, Walenz BP, Berlin K, Miller JR, Bergman NH and Phillippy AM. Canu: scalable and accurate long-read assembly via adaptive k-mer weighting and repeat separation. Genome Research. 2017;27 5:722–36. doi:10.1101/gr.215087.116.

14. Loman NJ, Quick J and Simpson JT. A complete bacterial genome assembled de novo using only nanopore sequencing data. Nat Methods. 2015;12 doi:10.1038/nmeth.3444.

15. Chin C-S, Peluso P, Sedlazeck FJ, Nattestad M, Concepcion GT, Clum A, et al. Phased diploid genome assembly with single-molecule real-time sequencing. Nature Methods. 2016;13:1050. doi:10.1038/nmeth.4035.

16. Michael TP, Jupe F, Bemm F, Motley ST, Sandoval JP, Loudet O, et al. High contiguity Arabidopsis thaliana genome assembly with a single nanopore flow cell. bioRxiv. 2017; doi:10.1101/149997.

17. Schloss PD, Jenior ML, Koumpouras CC, Westcott SL and Highlander SK. Sequencing 16S rRNA gene fragments using the PacBio SMRT DNA sequencing system. PeerJ. 2016;4:e1869. doi:10.7717/peerj.1869.

18. Benítez-Páez A, Portune KJ and Sanz Y. Species-level resolution of 16S rRNA gene amplicons sequenced through the MinION™ portable nanopore sequencer. GigaScience. 2016;5 1:4. doi:10.1186/s13742-016-0111-z.

19. Cusco A, Vines J, D’Andreano S, Riva F, Casellas J, Sanchez A, et al. Using MinION to characterize dog skin microbiota through full-length 16S rRNA gene sequencing approach. bioRxiv. 2017; doi:10.1101/167015.

20. Kerkhof LJ, Dillon KP, Häggblom MM and McGuinness LR. Profiling bacterial communities by MinION sequencing of ribosomal operons. Microbiome. 2017;5 1:116. doi:10.1186/s40168-017-0336-9.

21. Li C, Chng KR, Boey EJ, Ng AH, Wilm A and Nagarajan N. INC-Seq: accurate single molecule reads using nanopore sequencing. GigaScience. 2016;5:34. doi:D – NLM: PMC4970289 OTO – NOTNLM.

22. Benitez-Paez A and Sanz Y. Multi-locus and long amplicon sequencing approach to study microbial diversity at species level using the MinION portable nanopore sequencer. Gigascience. 2017;6 7:1–12.

23. Shin J, Lee S, Go M-J, Lee SY, Kim SC, Lee C-H, et al. Analysis of the mouse gut microbiome using full-length 16S rRNA amplicon sequencing. Scientific Reports. 2016;6:29681. doi:10.1038/srep29681.

24. Mitsuhashi S, Kryukov K, Nakagawa S, Takeuchi JS, Shiraishi Y, Asano K, et al. A portable system for rapid bacterial composition analysis using a nanopore-based sequencer and laptop computer. Scientific Reports. 2017;7 1:5657. doi:10.1038/s41598- 017–05772-5.

25. Quast C, Pruesse E, Yilmaz P, Gerken J, Schweer T, Yarza P, et al. The SILVA ribosomal RNA gene database project: Improved data processing and web-based tools. Nucleic Acids Research. 2013;41:590–6. doi:10.1093/nar/gks1219.

26. Singer E, Bushnell B, Coleman-Derr D, Bowman B, Bowers RM, Levy A, et al. High-resolution phylogenetic microbial community profiling. The Isme Journal. 2016;10:2020. doi:10.1038/ismej.2015.249.

27. Picher ÁJ, Budeus B, Wafzig O, Krüger C, García-Gómez S, Martínez-Jiménez MI, et al. TruePrime is a novel method for whole-genome amplification from single cells based on TthPrimPol. Nature Communications. 2016;7:13296. doi:10.1038/ncomms13296.

28. Rognes T, Flouri T, Nichols B, Quince C and Mahe F. Vsearch: a versatile open source tool for metagenomics. PeerJ. 2016;4 doi:10.7717/peerj.2584.

29. Schloss P, Westcott S, Ryabin T, Hall J, Hartmann M, Hollister E, et al. Introducing mothur: Open Source, Platform-independent, Community-supported Software for Describing and Comparing Microbial Communities. Appl Environ Microbiol. 2009; doi:10.1128/AEM.01541-09.

30. Westcott SL and Schloss PD. De novo clustering methods outperform reference-based methods for assigning 16S rRNA gene sequences to operational taxonomic units. PeerJ. 2015;3:e1487. doi:10.7717/peerj.1487.

31. Katoh K, Misawa K, Kuma K-i and Miyata T. MAFFT: a novel method for rapid multiple sequence alignment based on fast Fourier transform. Nucleic Acids Research. 2002;30 14:3059–66.

32. Katoh K and Toh H. Recent developments in the MAFFT multiple sequence alignment program. Briefings in Bioinformatics. 2008;9 4:286–98. doi:10.1093/bib/bbn013.

33. Pinto AJ and Raskin L. PCR Biases Distort Bacterial and Archaeal Community Structure in Pyrosequencing Datasets. PLoS ONE. 2012;7 8:e43093. doi:10.1371/journal.pone.0043093.

34. Jain M, Olsen HE, Paten B and Akeson M. The Oxford Nanopore MinION: delivery of nanopore sequencing to the genomics community. Genome Biology. 2016;17 1:239. doi:10.1186/s13059-016-1103-0.

35. Wang Q, Garrity G, Tiedje J and Cole J. Naive Bayesian classifier for rapid assignment of rRNA sequences into the new bacterial taxonomy. Appl Environ Microbiol. 2007;73 16:5261–7. doi:10.1128/AEM.00062-07.

36. Werner JJ, Koren O, Hugenholtz P, DeSantis TZ, Walters WA, Caporaso JG, et al. Impact of training sets on classification of high-throughput bacterial 16s rRNA gene surveys. ISME J. 2012;6 1:94–103. doi:10.1038/ismej.2011.82.

37. Feng W, Zhao S, Xue D, Song F, Li Z, Chen D, et al. Improving alignment accuracy on homopolymer regions for semiconductor-based sequencing technologies. BMC Genomics. 2016;17 7:521. doi:10.1186/s12864-016-2894-9.

38. RCoreTeam. R: A Language and Environment for Statistical Computing R Foundation for Statistical Computing, Vienna, Austria, 2014.

39. Edgar RC. MUSCLE: multiple sequence alignment with high accuracy and high throughput. Nucleic Acids Research. 2004;32 5:1792–7. doi:10.1093/nar/gkh340.

